# KRAS inhibition drives adaptive surfaceome remodeling and reveals ALPP/ALPPL2 as a targetable vulnerability in pancreatic cancer

**DOI:** 10.64898/2026.10.01.756025

**Authors:** Jonathan M. DeLiberty, Fan Yi Kong, Qijia Yu, Nicole Sindoni, Johnson Lin, Kalisa Kang, Elizabeth Y. Kim, Sabrina Chen, Aparna Padhye, Oluwadara Coker, Joshua H. Choe, Joao A. Paulo, Myungsun Shin, Lewis Hendrianto, Kazuki Takahashi, Jaclyn Varga-Wiles, Brendan D. Parent, Ziyue Li, Salome Shubitidze, Tobiloba E. Oni, David H. Peng, Timothy P. Heffernan, Shaojun Liu, Bryan K. S. Yeung, Bin Zou, Clint A. Stalnecker, Andrew J. Aguirre, Joseph D. Mancias

**Affiliations:** Department of Radiation Oncology, Dana-Farber Cancer Institute, Boston, MA; Department of Medical Oncology, Dana-Farber Cancer Institute, Boston, MA; Hale Family Center for Pancreatic Cancer Research, Dana-Farber Cancer Institute, Boston, MA; The Broad Institute of Harvard and MIT, Cambridge, MA; Harvard Medical School, Boston, MA; Lineberger Comprehensive Cancer Center, University of North Carolina at Chapel Hill, Chapel Hill, NC; Department of Cell Biology, Harvard Medical School, Boston, MA; Whitehead Institute for Biomedical Research, Cambridge, MA; David H. Koch Center for Integrative Cancer Research, Massachusetts Institute of Technology, Cambridge, MA; Therapeutics Discovery Division, University of Texas MD Anderson Cancer Center, Houston, TX USA; Axcynsis Therapeutics Pte. Ltd., Singapore; Department of Biochemistry and Biophysics, University of North Carolina at Chapel Hill, Chapel Hill, NC

**Keywords:** ALPP, ALPPL2, KRAS inhibition, RAS inhibition, PDAC

## Abstract

KRAS is mutated in over 90% of pancreatic ductal adenocarcinoma (PDAC) and is required for tumor initiation and maintenance. While clinical trials testing mutant-selective and pan-RAS inhibitors (KRASi) have demonstrated significant clinical benefit, genetic and non-genetic mechanisms of resistance limit the long-term efficacy of monotherapy strategies. Here, we used quantitative temporal analysis of the global, phosphorylation, and surface proteome to comprehensively identify non-genetic adaptive pathways and cell surface biomarkers in response to acute and long-term KRAS inhibition in PDAC. Through integrative machine learning analysis of the KRASi-proteome and publicly available datasets, we discovered top priority cell surface targets for combination therapy with KRASi. We identified Placental Alkaline Phosphatase (ALPP) as upregulated in response to KRASi through compensatory JAK2/STAT3 signaling. We developed a novel clinical grade ALPP/ALPPL2 (ALPG)-targeting antibody drug conjugate (ADC) with site-specific MMAE conjugation and demonstrated single agent efficacy in PDAC models, as well as enhanced internalization and anti-tumor efficacy in combination with KRASi. Together, these studies comprehensively define the KRASi-induced proteome and identify ALPP/ALPPL2 as a promising cell surface target for combination with KRASi to enhance response and forestall KRASi resistance.

## INTRODUCTION

Pancreatic ductal adenocarcinoma (PDAC) is the third leading cause of cancer deaths in the US, with a 5-year survival rate of approximately 8% (1). PDAC is characterized by mutations in *KRAS*, which activates oncogenic signaling through the mitogen-activated protein kinase (MAPK), phosphoinositide 3-kinase (PI3K), and other effector pathways (2,3). KRAS signaling is essential for tumor initiation and maintenance (4). Recent development of small molecule inhibitors of RAS proteins is transforming the treatment of multiple tumor types. Two mutant-specific KRAS^G12C^ inhibitors, adagrasib and sotorasib, have been approved for use in lung and colorectal cancer (5,6). Moreover, clinical trials of the RAS(ON) multi-selective inhibitor daraxonrasib have demonstrated a doubling of overall survival in patients with advanced PDAC and recent FDA approval (7–9) (NCT05379985, NCT06625320). Despite these advances, resistance to RAS inhibition is observed pre-clinically and in the clinic (10,11). Acquired resistance is frequently driven by genomic alterations, such as secondary KRAS mutations, gene amplifications, or mutations in downstream effectors like BRAF, which ultimately function to reactivate the mitogen-activated protein kinase (MAPK) pathway. Dual-targeting of these vertical pathways is currently being tested (12). Mechanistic dissection of treatment-induced resistance to KRAS inhibitor (KRASi) demonstrates that approximately 40% of patient resistance is driven by non-genetic mechanisms (13). These non-genetic resistance mechanisms are characterized, in part, by adaptive shifts in cell states and significant transcriptional remodeling, presenting opportunities for combination targeted strategies to therapeutically exploit these changes. We have previously used quantitative temporal proteomics to define adaptations to early pre-clinical KRAS^G12C^ inhibitors in PDAC and non-small cell lung cancer (NSCLC) (14) revealing rapid metabolic rewiring and cell cycle reactivation to bypass KRAS^G12C^ inhibition. This work informed combination therapies to increase the efficacy of KRAS^G12C^ inhibitors but focused on global proteomics, leaving cell surface adaptations largely unexplored.

We and others have demonstrated in preclinical models that combination of cytotoxic chemotherapies with KRASi can be an effective strategy to target the classical and basal transcriptional subtypes of PDAC (15); however, the toxicity of non-targeted systemic chemotherapy may limit combinations with more targeted agents. Antibody drug conjugates (ADCs) are an alternative approach to cytotoxic chemotherapy to deliver a cytotoxic payload to tumor cells. ADCs combine the specificity of monoclonal antibodies for their respective targets with the high potency of cytotoxic payloads via engineered linkers (16). While there are no current ADCs approved for PDAC, clinical successes in other cancer types provide a strong rationale for continued evaluation in PDAC and suggest a framework for defining the optimal characteristics of a candidate surface target (16). Oncogenic Kras signaling has been shown to functionally rewire the cell surfaceome in PDAC models to support cancer cell survival (17,18). In recent preclinical work, we have demonstrated that targeting the known cell surface proteins, MUC1 or ERBB2, with clinical ADC’s in combination with RAS inhibition can lead to enhanced efficacy over either agent alone (13,19). Thus, identifying additional novel cell surface proteins for therapeutic targeting may yield additional combination therapy strategies.

Here, we integrate global proteomics, phosphoproteomics, and a cell surface enrichment proteomic strategy to define the temporal evolution of the PDAC proteome with a specific focus on the ‘adaptive surfaceome’ during acute pharmacologic KRAS^G12D^ inhibition during response and resistance. We detect approximately 1,300 unique surface proteins that we prioritize using a machine learning algorithm to identify therapeutically tractable cell surface targets induced by KRASi in PDAC. Using this approach, placental alkaline phosphatase (ALPP), and its homolog ALPPL2 (ALPG), emerge as top KRASi-induced cell surface targets for ADC targeting in combination with (K)RAS inhibition in PDAC. ALPP is a glycosylphosphatidylinositol (GPI)-anchored alkaline phosphatase that catalyzes the hydrolysis of phosphoric monoesters (20) and is expressed primarily in the placenta with little to no expression in non-placental normal tissues (21,22). ALPP/ALPPL2 is inappropriately expressed in a variety of tumor types including ovarian cancer (23), gastric cancer (24), and PDAC (25) and has been targeted using ADC and cell-based therapy strategies in pre-clinical and early stage clinical studies (22,26,27). We demonstrate that KRASi-induced compensatory JAK2/STAT3 signaling upregulates ALPP/ALPPL2 and that targeting ALPP/ALPPL2 with a novel clinic-ready ADC in combination with pan-RAS(ON) inhibition drives deeper and sustained responses in PDAC models.

## RESULTS

### Temporal proteomic profiling reveals an adaptive rebound of essential cellular programs following KRAS inhibition

To characterize the proteomic adaptations driving the response and eventual resistance to KRAS inhibition (KRASi), we performed global quantitative temporal proteomics, phosphoproteomics, and surface-enriched proteomics (hereafter surfaceome) on a panel of ten PDAC cell lines treated with the KRAS^G12D^ inhibitor MRTX1133 (KRASi) (Fig. 1A, Supplementary Fig. S1A, Supplementary Table S1-2). We analyzed the acute response at 2 days across all ten lines. We further evaluated a subset of four lines at a sustained 7-day timepoint and at the point of acquired resistance (>2 months), where we found temporal rebound of phosphorylated ERK, consistent with previous reports (28) (Supplementary Fig. S1B-C). We also assessed treatment-naïve and treatment-resistant cell line-derived xenografts (CDXs) treated for 28 days with MRTX1133 to identify *in vivo* adaptations (Fig. 1A, Supplementary Fig. S1D).

**Figure 1.**
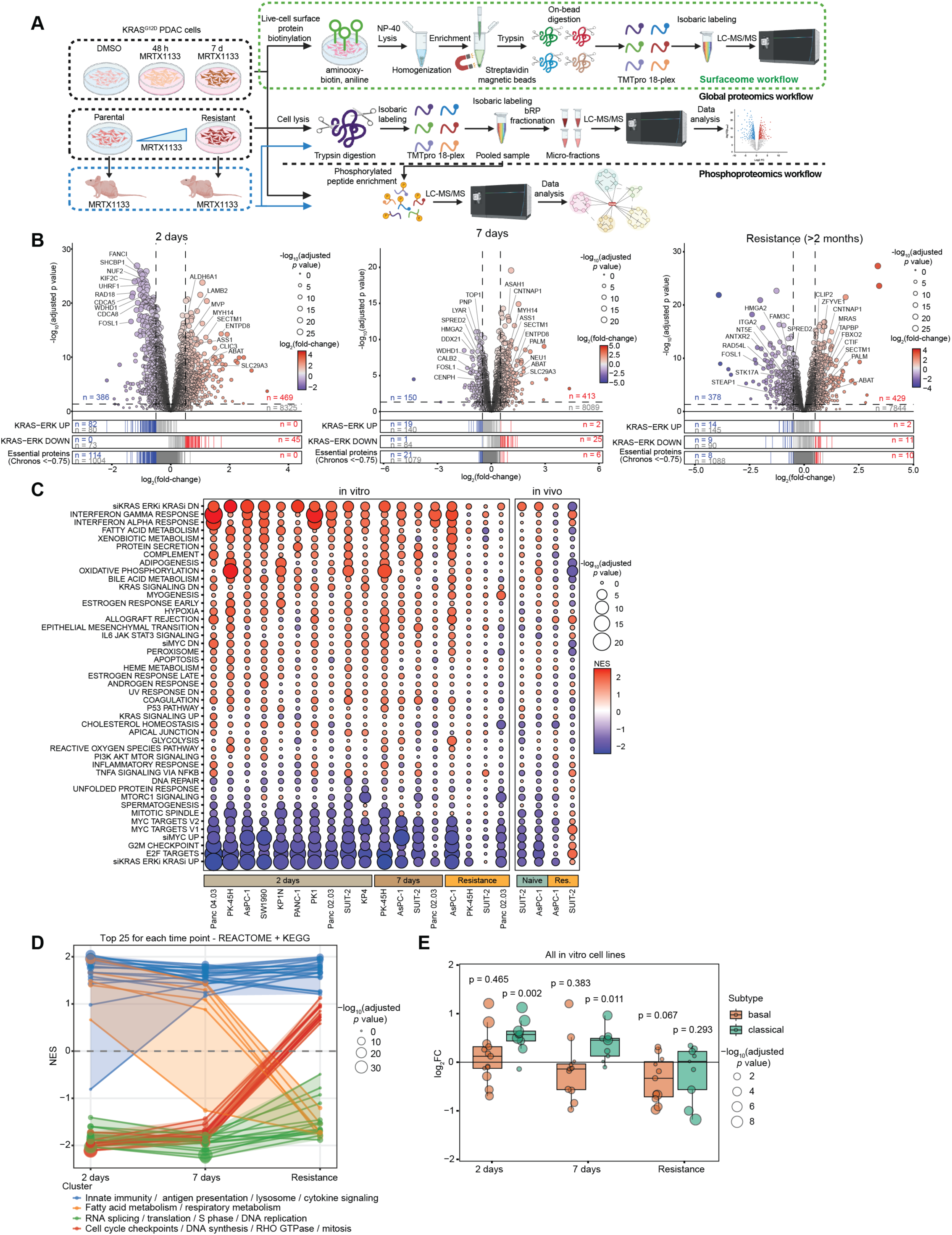
Temporal proteomic profiling maps the adaptive landscape of acquired resistance to KRAS^G12D^ inhibition in PDAC. **A.** Schematic workflow for determining the temporal KRAS-regulated proteome, phosphoproteome, and surfaceome (see text for further details). **B.** Aggregate volcano plots displaying relative protein abundance (log_2_ fold-change) versus significance (-log_10_ adjusted *p*-value) for KRASi-treated versus DMSO control samples. Analyses represent 10 PDAC cell lines at 2 days, and 4 cell lines longitudinally tracked to 7 days and acquired resistance. Core essential proteins (defined via DepMap) and KRAS-ERK regulated proteins (2) are plotted below, with significantly downregulated (adj. *p* < 0.05, log_2_(fold-change) < −0.5) in blue, significantly upregulated (log_2_(fold-change) > 0.5) in red, and non-significant in grey. **C.** Gene set enrichment analysis (GSEA) of Hallmark and KRAS-ERK (2) or MYC (29) signatures evaluated across in vitro (10 cell lines at 2 d, 4 cell lines at 7 d and resistance) and in vivo xenograft models (2 naïve and 2 with acquired resistance). **D.** Temporal GSEA trajectory mapping the net enrichment scores (NES) of the top 25 dysregulated REACTOME and KEGG pathways. Pathways were grouped into four distinct temporal clusters (k=4) utilizing agglomerative coefficient comparison and gap statistic analysis. Cluster keywords were consolidated using Szymkiewicz-Simpson overlap analysis (Supplementary Fig. S3A). **E.** Median log_2_(fold-change) of global proteins associated with established basal or classical PDAC transcriptional subtypes (72) across all three timepoints. Statistical significance was determined using a one-sample t-test comparing the mean log_2_ fold-change to 0.

Initial analysis revealed proteomic heterogeneity across the ten cell lines at day 2, a divergence that persisted in the four cell lines tracked until resistance, likely reflecting the known genetic and transcriptomic diversity across these cell lines (Supplementary Fig. S2A). Correlation analysis indicated that samples clustered primarily by cell line rather than by treatment duration (Supplementary Fig. S2B). To account for this baseline diversity, we incorporated a cell line-specific blocking term into our statistical model to account for inherent differences in baseline expression, allowing us to isolate the most conserved proteomic shifts. Consequently, we identified 855 proteins significantly differentially regulated at 2 days (adjusted *p* value < 0.05, |log_2_(fold-change)| > 0.5), 563 at 7 days, and 807 in resistant cells (Fig. 1B).

We observed strong suppression of proteins involved in KRAS-ERK signaling at 2 days of KRASi, which near-completely rebounded by 7 days and returned to baseline upon resistance (Fig. 1B). To determine if this signaling adaptation coincided with the recovery of critical survival programs, we cross-referenced our dataset with CRISPR essentiality scores for KRAS-mutant PDAC from the Cancer Dependency Map (DepMap). We observed a sharp downregulation of expression of 114 essential growth-driving proteins at day 2, with no corresponding upregulation of essential proteins (Fig. 1B). However, this anti-proliferative effect diminished over time: only 21 and 8 essential proteins remained suppressed at 7 days and at resistance, respectively (Fig. 1B). These trends were conserved across cell lines (Supplementary Fig. S2C), indicating a rapid signaling adaptation and normalization to KRASi occurring as early as 7 days.

Gene set enrichment analysis (GSEA) further defined this adaptive trajectory. Acute treatment suppressed established KRAS-dependent (2,3), MYC (29), and cell-cycle signatures (Fig. 1C, Supplementary Fig. S2D, Supplementary Table S3). While these pathways showed partial recovery by day 7, they largely reverted to baseline levels in resistant populations, a trend mirrored when comparing naïve versus resistant in vivo models. Despite widespread protein dysregulation at resistance (Fig. 1B), these alterations did not show strong enrichment patterns using Hallmark or KRAS-ERK/MYC pathways, reflecting a heterogenous and model-specific adaptive state.

To capture pathway dynamics, we mapped the net enrichment scores of the top 25 dysregulated KEGG and REACTOME pathways across all time points and identified four distinct temporal trajectories (Fig. 1D). Because many pathways share common genes, we applied the Szymkiewicz-Simpson overlap method to consolidate redundant terms (Supplementary Fig. S3A). First, innate immunity, antigen presentation, cytokine signaling, and lysosomal pathways remained enriched across all time points. This persistent lysosomal enrichment indicates that cells bypassing the KRASi-induced proliferative block adopt a distinct metabolic state, consistent with previous reports (30). Second, fatty acid and respiratory metabolism pathways were initially enriched at 2 days but became suppressed at resistance. Third, pathways driving RNA splicing, translation, S phase, and DNA replication were persistently suppressed. Finally, a fourth cluster comprising cell cycle checkpoints, DNA synthesis, RHO GTPases, and mitosis exhibited an adaptive rebound, transitioning from initial suppression to enrichment upon resistance. Collectively, we charted diverse signaling adaptations to KRASi that showed coordinated changes to metabolic pathways to support an adapted metabolic state, maintained suppression of distinct RNA splicing and biosynthetic pathways, and the rebound of cell cycle machinery to support cellular division and proliferation.

Finally, we examined whether KRASi adaptation involved shifts between established basal and classical PDAC transcriptional subtypes. Because these subtypes are characterized by empirically observed expression profiles rather than specific targetable biological pathways, we evaluated how KRASi impacted the global expression of their respective marker proteins. We found that KRASi did not drive coordinate changes in the expression of basal subtype proteins (Fig. 1E, Supplementary Fig. S3B). In contrast, acute KRAS inhibition induced a coordinated increase in classical protein expression. This aligns with previous reports demonstrating that KRAS suppression drives a cellular state shift toward a classical phenotype (31). However, we observed that this classical enrichment was transient; as cells adapted and acquired resistance, the expression of classical proteins subsided and largely returned to baseline levels.

### Temporal phosphoproteomics reveals dynamic rewiring of compensatory signaling networks to drive KRASi resistance

To delineate the dynamic signaling rewiring in response to KRAS inhibition, we quantified approximately 14,000 phosphosites across our temporal models. Correlation analysis demonstrated that samples clustered primarily by acute (2-day) versus sustained (7-day and resistance) KRASi treatment (Supplementary Fig. S4A, B). Using our cell line-blocked statistical model, we identified 3,552 phosphosites were significantly altered at 2 days, 1,402 at 7 days, and 1,762 upon resistance (Fig. 2A, Supplementary Table S4). Annotation of functionally validated regulatory phosphosites (32) revealed a progressive temporal shift, transitioning from profound initial downregulation to a rebound of upregulated sites driving compensatory pathway activation with sustained KRAS inhibition (Supplementary Fig. S4C).

**Figure 2.**
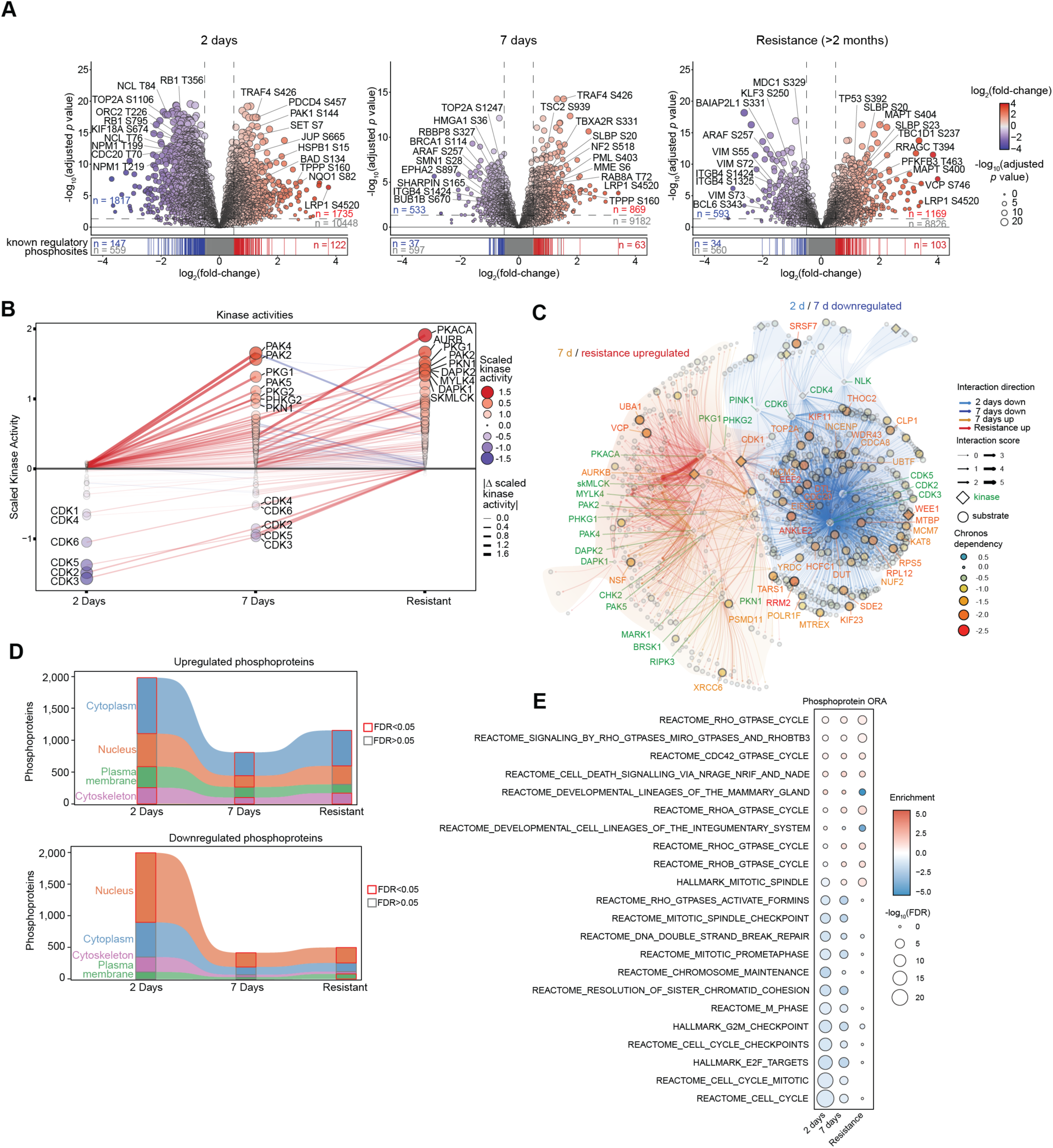
Temporal phosphoproteomics and kinome profiling reveal a spatially divergent adaptive signaling rebound driving KRASi resistance. **A.** Aggregate volcano plots displaying relative phosphosite abundance (log_2_ fold-change) versus significance (-log_10_ adjusted *p*-value) for KRASi-treated versus DMSO control samples. Analyses represent 10 PDAC cell lines at 2 days, and 4 lines longitudinally tracked to 7 days and acquired resistance. Functionally validated regulatory phosphosites (PhosphoSitePlus) are plotted below with significantly downregulated (adj. *p* < 0.05, log_2_ fold-change < −0.5) in blue, significantly upregulated (log_2_ fold-change > 0.5) in red, and non-significant in grey. **B.** Relative inferred activities for serine/threonine kinases across the temporal time course using kinase motif profiling. Line color denotes the direction of the activity, while width and transparency reflect the magnitude of change. **C.** Composite kinase-substrate networks mapping temporal adaptations. Node size and color denote genetic essentiality (DepMap Chronos scores), with kinases represented as diamonds and substrates as circles. Edge color indicates the direction of association (up- or downregulated phosphosites), and edge width reflects the magnitude of the interaction score. Top-quantile kinases (green) and highly essential proteins are labeled. **D.** Subcellular compartment distribution of significantly up- (top) and down-regulated (bottom) phosphoproteins. Significantly enriched compartments (determined by one-tailed hypergeometric testing) are outlined in red. **E.** Overrepresentation analysis (ORA) of significantly dysregulated phosphoproteins against KEGG, REACTOME, and HALLMARK gene sets. Statistical enrichment was determined via one-tailed hypergeometric testing at each timepoint.

Functionally, these regulatory sites highlighted two distinct adaptive phases. Acute KRASi severely suppressed nodes governing cell-cycle progression and DNA replication (e.g. RB1, TOP2A, BRCA1, CDC20) while transiently upregulating early stress-adaptive and metabolic responses (e.g., PDCD4, TSC2), cytoskeletal remodeling (e.g., PAK1, ITGB4) and signaling rewiring (e.g., EPHA2, LRP1). In contrast, resistant cells bypassed this initial proliferative block by reestablishing cytoskeletal dynamics, vesicle trafficking, and metabolic signaling (e.g., MAPT, TP53, PFKFB3, mTOR/Rag regulators). Ultimately, sustained KRAS inhibition leads to signaling adaptations that allow cells to resume outgrowth.

To deduce the specific kinases driving these changes, we employed kinase motif profiling (33). Correlation analysis of kinase activities showed high agreement across timepoints (Supplementary Fig. S4D). Acute KRAS inhibition significantly suppressed cell cycle regulatory CDKs, which progressively rebounded over time as cells regained their proliferative capacity (Fig. 2B, Supplementary Fig. S4E, Supplemental Table S5). Concurrently, sustained KRAS inhibition induced a strong compensatory upregulation of kinases governing cytoskeletal dynamics (PAKs, PKNs, MYLK, SKMLCK), cell death responses (DAPKs), and mitotic entry (AURKB).

To map these dynamic kinase activities to their functional effectors, we constructed kinase-substrate networks (3) and integrated genetic dependency scores from the Cancer Dependency Map (DepMap) (Fig. 2C, Supplementary Fig. S4F–H). At 2 days, we observed a massive enrichment of downregulated interactions involving essential proteins, reflecting the initial inhibition of proliferation. However, sustained KRAS inhibition yielded very few upregulated sites on essential proteins, and initially suppressed essential networks rebounded over time. This suggests that acquired resistance is driven primarily by the reactivation of initially suppressed essential pathways rather than the stimulation of orthogonal, *de novo* essential networks.

Finally, we mapped the subcellular compartments and pathways underlying these adaptations (Fig. 2D, E). Downregulated phosphosites were predominantly nuclear, reflecting an acute halt in cell cycle regulatory proteins that rebounded upon resistance. Conversely, upregulated phosphosites were at the plasma membrane and within the cytoskeleton and cytoplasm, highlighted by the enrichment of upregulated phosphoproteins in RHO GTPase pathways (CDC42, RHOA/B/C). This spatial divergence demonstrates that PDAC cells survive the initial KRASi-induced proliferative block by engaging pro-survival, motility, and EMT-associated signaling within cytoplasmic and plasma membrane compartments. Crucially, this enrichment of adaptive signaling at the plasma membrane suggested that the cell surface undergoes coordinated structural remodeling during KRASi, prompting us to investigate the KRAS-regulated surfaceome.

### Sustained KRAS inhibition induces robust remodeling of the cell surfaceome

Our global and phosphoproteomics analyses defined the temporal evolution of adaptive intracellular pathways following KRAS inhibition. Because this reprogramming integrated heavily with the cellular architecture and plasma membrane, we sought to determine if the cell surface undergoes coordinated structural alterations that might be exploited therapeutically with antibody-drug conjugates (ADCs). To selectively profile this compartment, we employed a live-cell aminooxy-biotin (BAMX)-based labeling strategy followed by streptavidin magnetic bead enrichment and mass spectrometry-based proteomics (Fig. 1A) (34). Using an annotated list of empirically determined and predicted surface proteins (35–37), we detected over 900 unique surface proteins during acute and sustained KRASi treatment, and over 800 in the acquired resistance models (Fig. 3A, Supplementary Fig. S5A, Supplementary Table S6). We observed profound dysregulation across all three timepoints, including altered expression of established clinical targets such as MUC1, SLC44A4, CEACAM5, and FGFR3.

**Figure 3.**
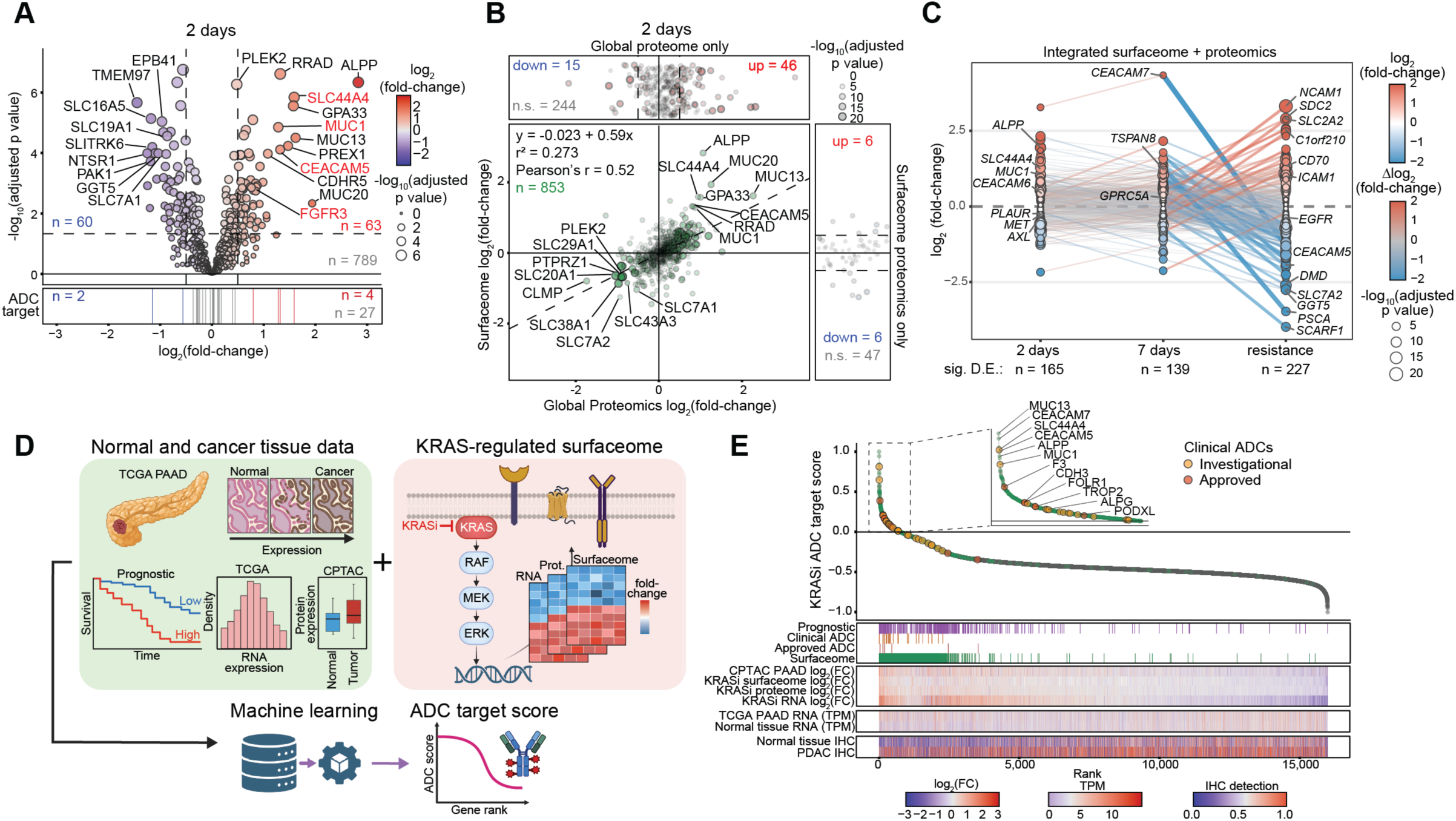
Integration of temporal surfaceomics and machine learning identifies ADC targets in KRAS-inhibited PDAC. **A.** Aggregate volcano plot showing differential expression (log_2_ fold-change) versus significance (-log_10_ adjusted *p*-value) of cell surface proteins quantified by BAMX surface protein enrichment in 10 PDAC cell lines treated with KRASi for 2 days. Clinically investigated ADC targets are annotated below, with significantly upregulated (adj. *p* < 0.05, log_2_ fold-change > 0.5) highlighted in red and downregulated (adj. *p* < 0.05, log_2_ fold-change < −0.5) in blue. **B.** Correlation analysis comparing surface-annotated proteins detected by global proteomics versus BAMX surface enrichment. Green dots indicate proteins shared between modalities, while red (top) and blue (right) dots represent proteins uniquely detected by global proteomics or BAMX enrichment, respectively. Significance thresholds (adj. p < 0.05, |log2FC| > 0.5) are denoted for uniquely detected proteins. **C.** Temporal trajectory map of surface protein expression (log_2_FC) derived from the integrated proteomics and surfaceome datasets. Line color and thickness indicate the magnitude of expression change (Δlog_2_FC) between adjacent timepoints. **D.** Schematic of the multi-omic machine learning pipeline used to compute the KRASi ADC Target Score. The ridge logistic regression model was trained using healthy tissue profiles, PDAC tumor expression/survival data (TCGA, CPTAC, Human Protein Atlas), and KRAS-regulated dynamics from published transcriptomics and our integrated in-house surfaceomics. **E.** Rank plot of the computed KRASi ADC Target Scores for all detected genes, derived by log-odds linear prediction from the trained model. Targets with clinically investigated (yellow) or FDA-approved (orange) ADCs are highlighted. The multi-omic feature heatmaps utilized for training are aligned below. The inset magnifies the top-scoring candidates (score > 0), highlighting priority targets for ADC development.

To expand our coverage of potential ADC targets, we integrated our surface-enriched proteomics with our global proteomic datasets. The two modalities exhibited high quantitative agreement, sharing over 800 surface proteins. Global proteomics captured an additional 300 surface annotated proteins, while fewer than 100 were unique to the BAMX enrichment (Fig. 3B, Supplementary Fig. S5B, C). By combining these datasets using Stouffer’s weighted z-score, we mapped a highly dynamic, integrated KRAS-regulated surfaceome (Fig. 3C, Supplemental Table S7). This integration revealed an expanded surfaceome with 165 differentially regulated surface proteins at 2 days, 139 at 7 days, and 227 upon acquired resistance. Additionally, this integration significantly broadened our pool of candidate ADC targets (Supplementary Fig. S5D). While surfaceome alterations were largely conserved between the 2-day and 7-day timepoints, the landscape diverged dramatically at resistance. This suggests that surface markers upregulated during the initial response to KRASi may not be sustained once cells fully evade treatment.

Having mapped the KRAS-regulated surfaceome, we engineered a machine learning (ML) pipeline to systematically prioritize ADC targets. Conceptually, an ideal therapeutic target is characterized by low expression in healthy tissues, high baseline expression in PDAC, and a robust induction upon KRAS suppression. The model incorporated multi-omic features, including healthy tissue expression databases from the Human Protein Atlas (38), normal and cancer immunohistochemistry (IHC) scores (39), published MRTX1133-driven transcriptomics (40), and our newly defined KRAS-regulated proteomic datasets (Fig. 3D). Because our goal was to identify targets suitable for early combination therapy, and we observed profound reprogramming of the KRAS-regulated surfaceome at resistance, we restricted our training set to the acute and sustained response timepoints. We identified 32 surface proteins with clinically investigated ADCs that were upregulated at 2 or 7 days of KRASi to identify our training set. We used 10 molecular features to train a ridge logistic regression model to compute a “KRASi ADC target score”, which prioritizes features that were shared across the training set (Fig. 3E, Supplemental Table S8). This model was associated with positive coefficients for surface-protein annotation, KRASi-induced expression and expression in cancer tissues, and negative coefficients for normal tissue expression. Validating this approach, we found established clinical ADC targets such as SLC44A4 (CTL4), CEACAM5 (CEA), and MUC1 among our highest hits.

Analyzing the top-scoring hits as targets for ADCs confirmed they predominantly exhibit strong RNA and protein upregulation by KRASi, low normal tissue expression, and high baseline prevalence in PDAC tumors (Supplementary Fig. S5F). Furthermore, we benchmarked these top hits against low-scoring targets and known resistance-associated surface receptors (e.g. MET, EGFR, AXL). Low-scoring hits were predominantly characterized as not being upregulated by KRAS inhibition, lower PDAC tumor expression, or not surface targets. Ultimately, this pipeline translates the dynamic surfaceome reprogramming induced by KRAS inhibition into a robust, machine-learned framework for prioritizing tractable ADC combinations.

### ALPP is a therapeutically tractable surface protein upregulated by KRASi

We evaluated the top candidates from the machine learning approach and found targets that were increased across our panel of cell lines (e.g. ALPP, THBS1, ICAM1), whereas others were specific to one or a few cell lines (e.g. MUC13, CEACAM7, SLC44A4) (Supplementary Fig. S5E). We selected several clinical and pre-clinical surface targets of interest to validate by flow cytometry including ALPP, ICAM1, CEACAM5, and *TACTSD2*/TROP2 (Fig. 4A-B, Supplementary Fig. S6A). We found increased surface expression by flow cytometry following KRASi across the selected candidates, validating this proteomic and bioinformatic strategy can robustly identify and prioritize therapeutic targets.

**Figure 4.**
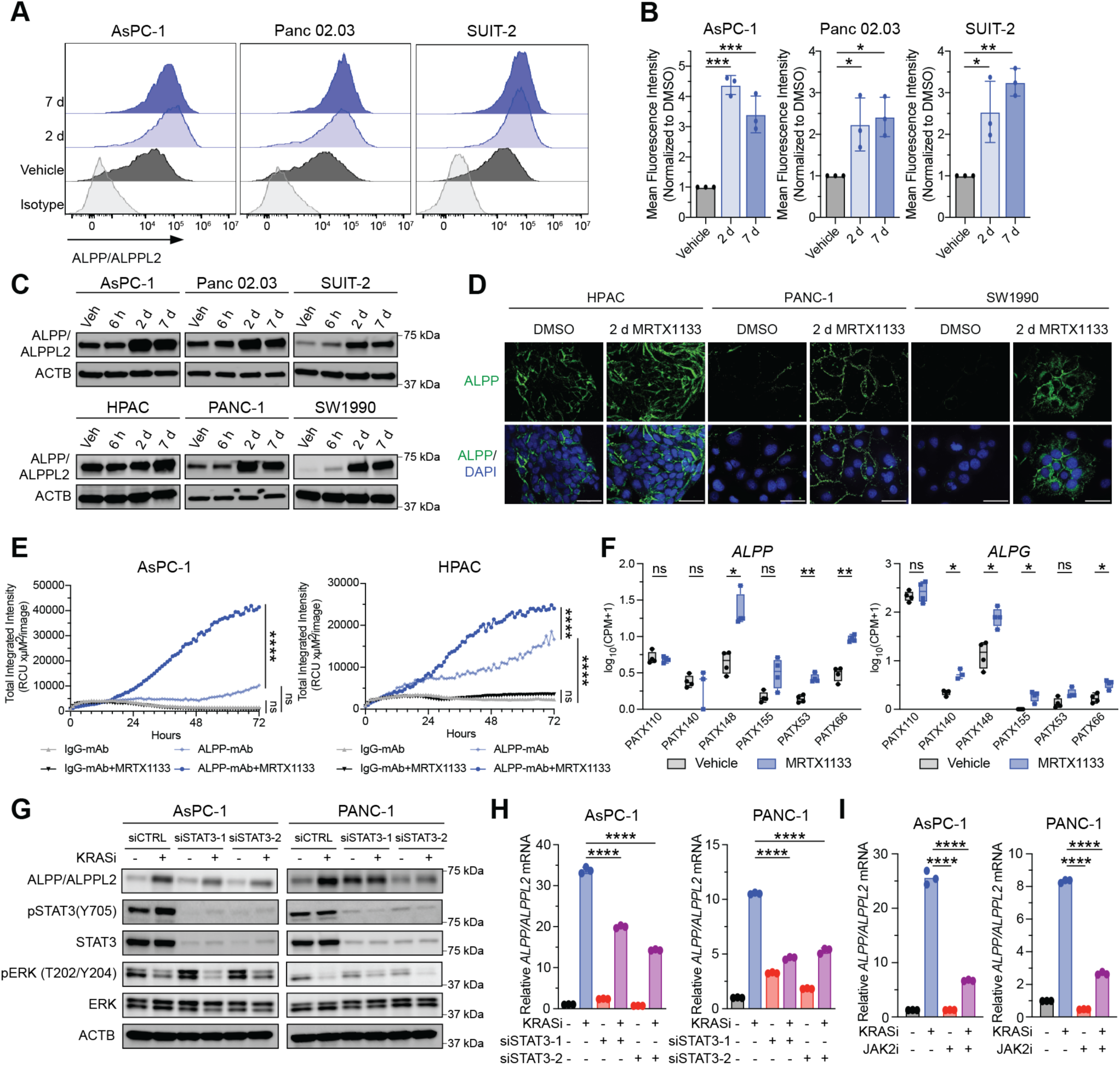
ALPP/ALPPL2 surface expression is upregulated by KRASi via a JAK2/STAT3 mechanism. **A**. Flow cytometry analysis of ALPP/ALPPL2 expression. PDAC cells were treated with Vehicle, KRASi for 2 days (MRTX1133, 100 nM), or KRASi for 7 days (MRTX1133, 20 nM). Representative histograms of three independent experiments are shown. **B**. Quantification of ALPP/ALPPL2 surface expression from flow cytometry in **A**. Mean fluorescence intensity was normalized to vehicle control; error bars denote mean ± s.d., n = 3 biological replicates. Statistical significance was determined by one-way ANOVA. **C**. ALPP/ALPPL2 protein levels in PDAC cells treated for 6 hours or 2 days with KRASi (MRTX1133, 100 nM for AsPC-1, Panc 02.03, SUIT-2, HPAC, and SW1990; 1 µM for PANC-1) or for 7 days with KRASi (MRTX1133, 20 nM for AsPC-1, Panc 02.03, SUIT-2, HPAC, and SW1990; 1 µM for PANC-1) determined by western blotting. **D**. Immunofluorescence analysis of PDAC cells stained for ALPP treated with DMSO or 100 nM MRTX1133 for 2 days. Scale bar = 50 µm. **E.** Internalization of the anti-ALPP/ALPPL2 or IgG antibody (10 nM), quantified by total integrated density by Incucyte live-cell imaging. AsPC-1 and HPAC cells were treated with and without KRASi (MRTX1133; 100 nM for AsPC-1, 10 nM for HPAC). **F.** Published RNAseq data from six KRAS^G12D^ PDAC PDX models treated with 30 mg/kg BID MRTX1133 for 3 days (31). Boxes show median and interquartile range with individual tumors overlaid. Unpaired t-tests on log_10_(CPM+1) values were performed within each model, Šídák-Bonferroni corrected for multiple comparisons (ns, not significant; *P<0.05; **P<0.01). **G.** ALPP/ALPPL2, phospho-STAT3 (Y705), STAT3, phospho-ERK (T202/Y204), and ERK protein levels in PDAC cells treated with a non-targeting control siRNA (siCTRL) or two different siRNAs targeting *STAT3* (siSTAT3) for 2 days before treatment with KRASi (100 nM MRTX1133 for AsPC-1 and SW1990, 1 µM MRTX1133 for PANC-1) for 2 days, determined by western blotting. Representative of three biological replicates. **H**. Relative *ALPP/ALPPL2* mRNA levels measured by qPCR in the same samples as shown in **G**, error bars denote mean ± s.d., n = 3 independent repeats. Statistical significance was determined by one-way ANOVA. Representative of three biological replicates. **I**. Relative *ALPP/ALPPL2* mRNA levels in PDAC cells treated with DMSO, KRASi (MRTX1133, 100 nM for AsPC-1; 1 µM for PANC-1), JAKi (Cucurbitacin I, 250 nM), or a combination of KRASi and JAKi for 2 days as measured by qPCR, error bars denote mean ± s.d., n = 3 independent repeats; representative of 3 biological replicates. Statistical significance was determined by one-way ANOVA

We focused our subsequent work on alkaline phosphatase proteins, placental (ALPP) and its close homolog ALPPL2 (ALPG), as these were the most consistently upregulated surface protein across cell lines and validated by flow cytometry. ALPP and ALPPL2 are membrane-bound glycosylphosphatidylinositol (GPI)-anchored phosphatases that catalyze the hydrolysis of phosphate esters (20). ALPP/ALPPL2 is normally expressed in the placenta with reports suggesting it facilitates nutrient transport (21,41), but it has also been found to be aberrantly overexpressed in a number of different cancer types, including ovarian cancer, lung cancer, and PDAC. Indeed, the Genotype-Tissue Expression (GTEx) Portal demonstrates low ALPP expression in non-diseased tissue sites (Supplementary Fig. S6B). We found temporal induction of ALPP/ALPPL2 expression following KRASi in a panel of PDAC cell lines by immunoblotting (Fig. 4C). Importantly, we observed increased cell-surface expression by 2 days of KRAS inhibition using immunofluorescence (Fig. 4D). A critical determinant of ADC efficacy is cell surface protein-antibody internalization, which recent clinical biomarker data suggests may be more predictive of ADC response as opposed to surface expression alone. More specifically, a TROP2-targeting ADC clinical trial in NSCLC patients (NCT04526691) observed that treatment response correlated with a lower membrane-to-cytoplasm ratio (indicative of greater TROP2 cycling/internalization), instead of total TROP2 expression by conventional IHC. Therefore, we evaluated the level of ALPP/ALPPL2 internalization in response to KRAS inhibition. We identified a significant increase in the internalization rate of ALPP surface protein upon KRASi, in line with the increase in cell surface ALPP expression (Fig. 4E, Supplementary Fig. S6C).

Analysis of published KRASi RNA-seq datasets identified that ALPP and ALPG are transcriptionally upregulated following KRASi treatment in both *in vitro* and *in vivo* models of PDAC, including mice harboring patient-derived xenograft (PDX) PDAC tumors (Fig. 4F, Supplementary Fig. S6D) (31,40), suggesting ALPP and ALPG are targets of a KRAS-dependent transcriptional response. Signal transducer and activator of transcription 3 (STAT3), a critical transcription factor that integrates diverse oncogenic signals to promote tumorigenesis and anti-apoptotic activities (42), has been suggested as a putative regulator of ALPP in response to EGFR inhibition in EGFR-mutant non-small cell lung cancer (43). In the context of KRAS-mutant PDAC, KRAS inhibition results in upregulated JAK signaling and increased active phospho-STAT3 (44,45). Taken together, these findings suggest that ALPP and ALPG may be upregulated following KRAS inhibition by a STAT3-dependent transcriptional mechanism. To evaluate this, we utilized genetic knockdown of STAT3 and pharmacologic targeting of JAK2/STAT3 in our PDAC models. However, RNAi-mediated STAT3 ablation decreased KRASi-induced upregulation of ALPP protein expression and ALPP mRNA levels (Fig. 4G, H). Similarly pharmacological inhibition of the JAK2/STAT3 pathway using Cucurbitacin I (JAKi) abrogated the upregulation of ALPP/ALPPL2 mRNA levels induced by KRAS inhibition (Fig. 4I). Taken together, ALPP surface protein upregulation in response to KRASi treatment is mediated through a STAT3-dependent transcriptional mechanism.

Cell surface proteins upregulated in response to KRASi have been implicated as mediators of KRASi resistance, including EGFR and ERBB3, primarily by the reactivation of RTK signaling. We evaluated whether ALPP/ALPPL2 upregulation confers resistance to KRAS inhibition. RNAi-mediated knockdown of ALPP/ALPPL2 in PDAC cell lines did not significantly alter PDAC cell line sensitivity to MRTX1133 treatment (Supplementary Fig. S6E, F).

To take advantage of the KRASi-induced upregulation and increased internalization of ALPP/ALPPL2 for therapeutic targeting, we developed a next-generation ALPP/ALPP2 targeting ADC and evaluated it in combination with KRASi.

### AT2604 is an ADC targeting ALPP/ALPPL2 and mediates anti-tumor efficacy in preclinical models

Axcynsis Therapeutics has developed a next-generation anti-ALPP/ALPPL2 ADC (AT2604) through the engineering of a murine monoclonal antibody recognizing human ALPP (clone H17E2; NCT05229900). Beginning with the variable region of the original murine anti-ALPP antibody (46), the complementarity determining regions (CDRs) were grafted onto the closest human germline framework identified by the IMGT/V-QUEST system (47). An increase in binding affinity over AT04-5A (clone H17E2) was observed upon reintroducing Glu56 to the framework sequence that modelling predicted might reestablish a key interaction in the framework region, resulting in AT04-13A (Fig. 5A). Next, a sequence analysis identified an Asp54-Gly55 (DG) motif as an aspartate isomerization hotspot that was ameliorated through a G55Q mutation (48). Engineered cysteines Ala118Cys and Ser239Cys were introduced for site-specific bioconjugation to provide AT04-23A, which maintained strong affinity and selectivity over the other human alkaline phosphatase isozymes (ALPI and ALPL) and cross-reactivity to cynomolgus ALPPL2 but no reactivity to murine ALPPL2 (22) (Fig. 5A-B). In a final optimization step to abrogate antibody effector function, Fc-silencing LALA (Leu234Ala/Leu235Ala) mutations were introduced to provide AT04-26A (Supplementary Fig. S7A) (49). Antibody AT04-26A was then bioconjugated with a clinically validated mc-vc-PAB linker and monomethyl auristatin E (MMAE) payload (50) to provide the final ADC, termed AT2604, with a highly controlled DAR4 (Supplementary Fig. S7B).

**Figure 5.**
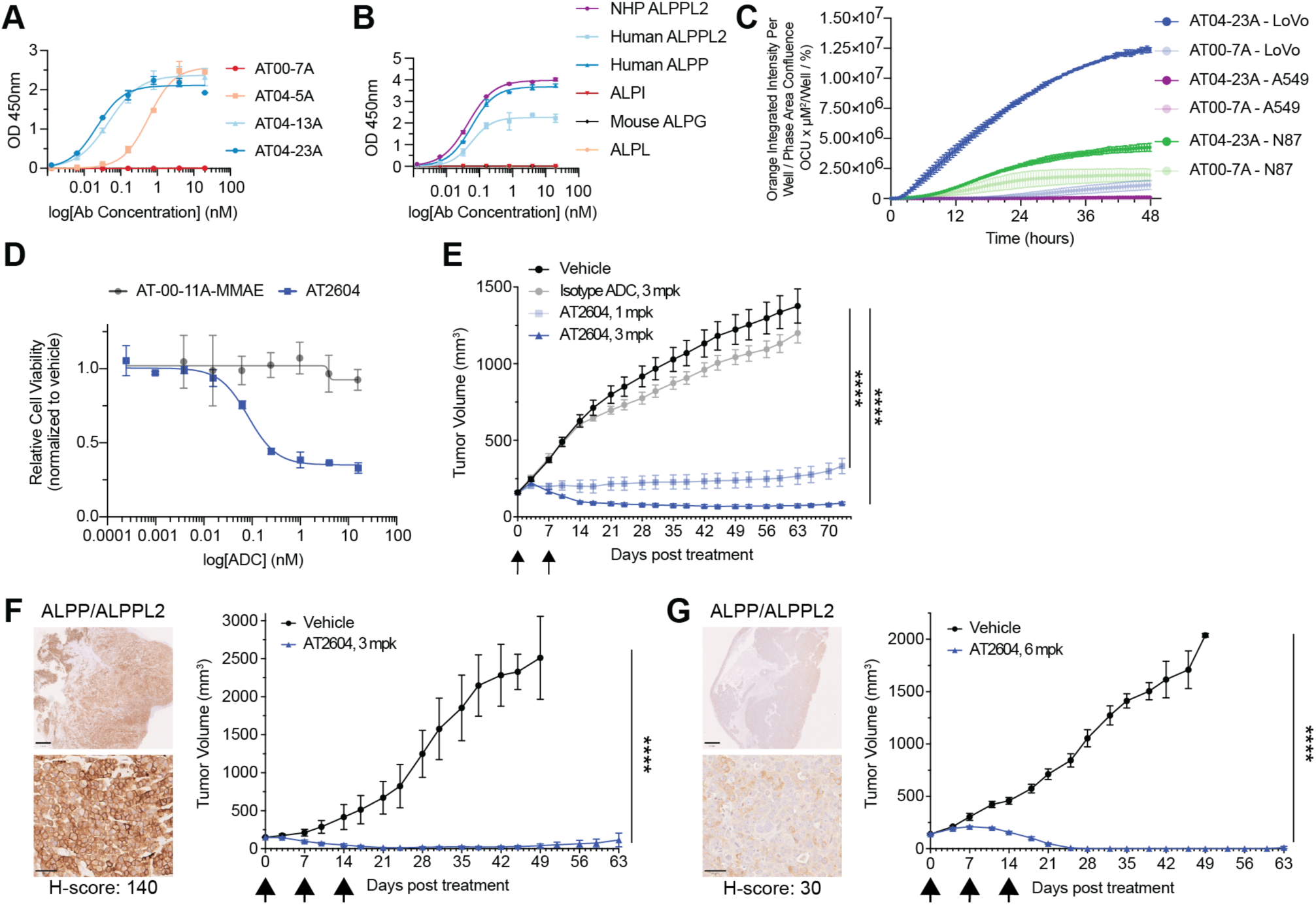
Development and preclinical activity of AT2604, an ADC targeting ALPP/ALPPL2. **A.** Binding affinity of AT00-7A, AT04-5A, AT04-13A, and AT04-23A to human ALPPL2 by ELISA assay. **B.** Antigen selectivity ELISA assay of optimized antibody AT04-23A for human ALPP, human ALPPL2, other human alkaline phosphatases (ALPLI, ALPL), and ALPPL2 from non-human species. **C.** Internalization of AT04-23A (2 µg/mL) and isotype control (AT00-7A, 2 µg/mL) in NCI-N87, LoVo, and A549 cells over 48 hours assessed by Incucyte live cell imaging. **D.** Anti-proliferative cytotoxicity of AT2604 relative to the isotype ADC (AT-00-11A-MMAE) in NCI-N87 cells. **E.** Balb/c nude mice harbouring subcutaneous NCI-N87 cell line-derived tumors were treated with vehicle or indicated doses of isotype ADC or AT2604 IV QWx2 (n=6 mice per arm), and tumor volumes were monitored by caliper measurement. **F.** IHC images of GA2419 PDX tumor stained with anti-ALPP/ALPPL2 antibody and corresponding H-score. Top image scale bar = 1 mm; bottom image scale bar = 50 µm (left). Tumor volume growth curve of AT2604 (3 mpk IV QWx3) treated patient-derived xenograft (PDX) model of gastric cancer, GA2419, n=3 mice per arm (right). **G.** IHC images of GA6203 PDX tumor stained with anti-ALPP/ALPPL2 antibody and corresponding H-score. Top image scale bar = 1 mm; bottom image scale bar = 50 µm (left). Tumor volume growth curve of AT2604 treatment (6 mpk IV QWx3) in a PDX model of gastroesophageal junction cancer (GEJ), GA6203, n=3 mice per arm, (right). Statistical analysis was performed using one-way ANOVA followed by Tukey’s multiple comparisons test. ****P<0.0001.

Anti-ALPP/ALPPL2 antibody AT04-23A demonstrates robust internalization relative to the isotype control (AT00-7A) in high ALPP/ALPPL2 expressing gastric (NCI-N87), and colon (LoVo) cancer cell lines but not in the ALPP/ALPPL2-negative A549 non-small cell lung cancer line (Fig. 5C). Concordant with robust internalization in NCI-N87 cells, AT2604 conferred a significant anti-proliferative effective relative to the isotype ADC in a cell culture-based cell viability assay (Fig. 5D). Cell-derived xenografts (CDX) of NCI-N87 treated with AT2604 demonstrated deep and sustained tumor regressions in comparison to vehicle or Isotype ADC administration consistent with potent in vivo activity of AT2604 in a high-ALPP/ALPPL2 expressing model (Fig. 5E). We next evaluated AT2604 in PDX models with varying baseline levels of ALPP/ALPPL2. AT2604 administration in a ALPP/ALPPL2-high expressing gastric cancer PDX model, GA2419 (H-score: 140; 3 mg/kg QWx3) achieved >90% tumor regression (Fig. 5F). Notably, AT2604 treatment in an ALPP/ALPPL2-low expressing gastroesophageal junction cancer (GEJ) PDX model, GA6203 (H-score: 30; 6 mg/kg QWx3), led to a sustained complete response highlighting the efficacy of AT604 in both low and high-expressing ALPP/ALPPL2 tumor models (Fig. 5G). AT2604 administration was well tolerated in these in vivo models with no weight loss observed in the treatments group (Supplementary Fig. SC-E) likely reflecting a tolerable level of free payload and low non-specific antibody uptake given AT2604 does not recognize murine ALPP/ALPG. Taken together, we find that AT2604 is a novel ALPP/ALPPL2-targeting ADC with high specificity and robust efficacy in a variety ALPP/ALPPL2-expressing cancer types.

### Combination of RAS inhibitor and anti-ALPP/ALPPL2 ADC is an efficacious therapeutic strategy in PDAC

AT2604 presents a potential clinic-ready ADC to therapeutically exploit the increase of surface ALPP/ALPPL2 upon KRAS inhibition in KRAS-mutant cancers, particularly PDAC. To determine whether this ADC could bind to the surface of our panel of PDAC cell lines, we utilized the parent antibody (AT04-26A). PDAC cell lines demonstrated robust binding of the ALPP/ALPPL2 parent antibody relative to IgG control (Fig. 6A, Supplementary Fig. S7F). We also observed a significant increase in surface expression of ALPP/ALPPL2 following 2 days or 7 days of KRASi treatment in the majority of cell lines tested, except for the ALPP/ALPPL2-negative PK-45H cell line (Fig. 6A, Supplementary Fig. S7F). We next sought to determine if ALPP/ALPPL2 upregulation in response to KRASi was a phenomenon limited to PDAC. We evaluated multiple KRAS-mutant ovarian cancer cell lines and observed robust upregulation of ALPP/ALPPL2 surface expression following KRASi/RASi treatment (Supplementary Fig. S7G), suggesting this phenotype can be extended to other KRAS-mutant cancer types. We once again observed that this upregulation is mediated by a STAT3-dependent transcriptional mechanism in KRAS-mutant ovarian and lung cancer cell lines (Supplementary Fig. S7H). We then sought to determine if the anti-ALPP/ALPPL2 ADC, AT2604, could be internalized effectively by our PDAC cells. We demonstrate that AT2604 was internalized to a greater extent than its parent antibody (AT04-26A), and a significantly greater amount than the isotype control ADC (AT-00-11A-MMAE) (Fig. 6B, Supplementary Fig. S8A). Interestingly, the addition of KRASi treatment increased the internalization of the parent antibody and AT2604 but not the isotype control in all cell lines tested (Fig. 6B, Supplementary Fig. S8A). This is likely due, in part, to the increase in ALPP/ALPPL2 surface expression upon KRASi treatment.

**Figure 6.**
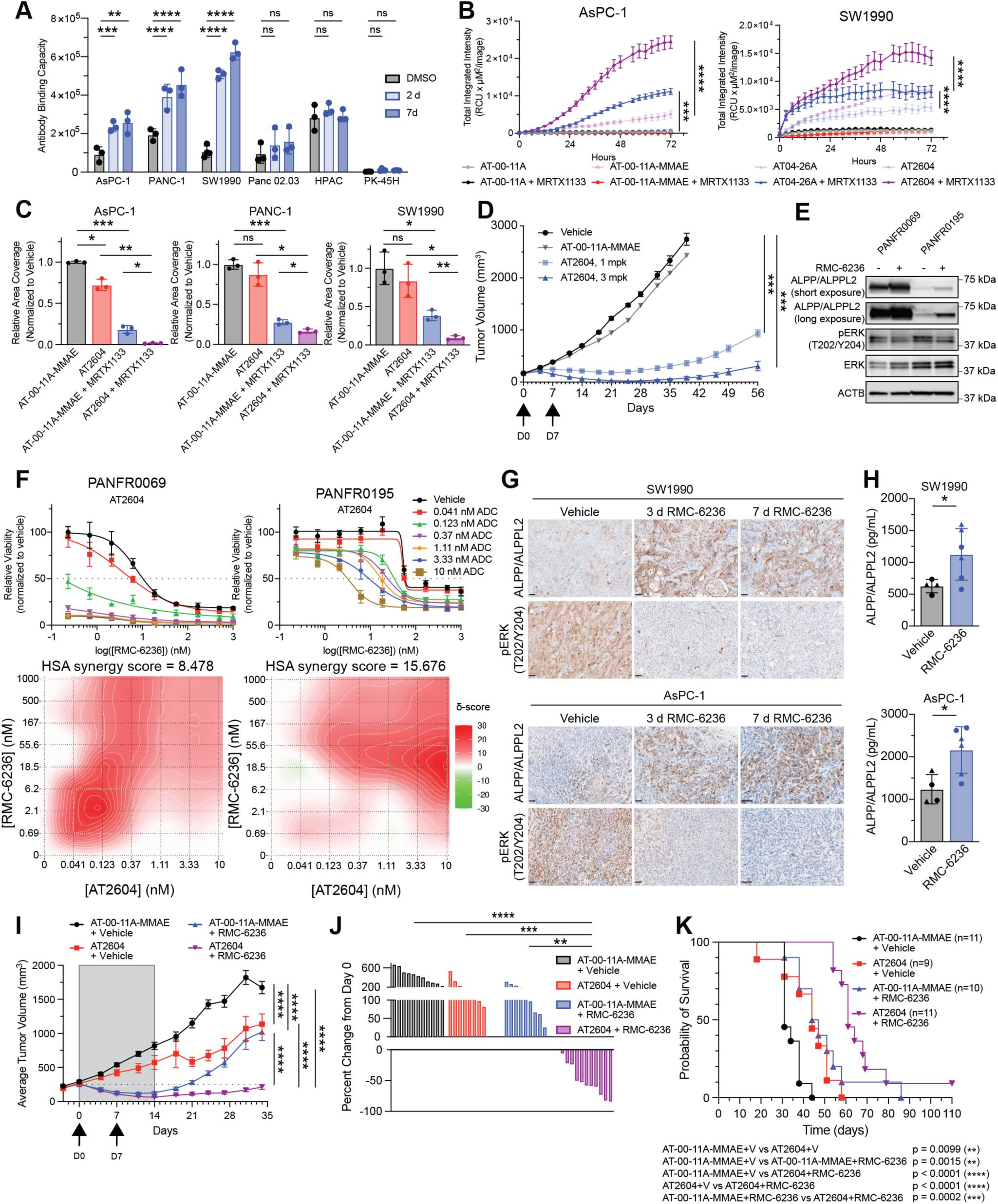
Combination of RASi and ALPP/ALPPL2 ADC leads to sustained tumor growth inhibition in PDAC models. **A.** Antibody-binding capacity of ALPP/ALPPL2 in PDAC cells treated with DMSO, KRASi for 2 days (MRTX1133; 100 nM for AsPC-1, Panc0203, HPAC, SW1990, and PK-45H; 1 µM for PANC-1), or KRASi for 7 days (MRTX1133; 20 nM for AsPC-1, Panc0203, HPAC, SW1990, and PK-45H; 1 µM for PANC-1) as measured by flow cytometry. Each data point represents an independent experiment, error bars denote mean ± s.d., n = 3 independent repeats. Statistical significance was determined using two-way ANOVA followed by Tukey’s multiple comparisons test. **B.** Internalization of the indicated antibody or antibody-drug conjugate (10 nM) with or without KRASi (MRTX1133, 100 nM for AsPC-1 and SW1990) was quantified by total integrated density by Incucyte live-cell imaging. Internalization curves represent three technical replicates; error bars denote mean ± s.e.m, representative of three biological replicates. Statistical significance was determined by one-way ANOVA followed by Tukey’s multiple comparisons test. **C.** Quantification of relative area coverage from crystal violet assays normalized to IgG-ADC control (AT-00-11A-MMAE). PDAC cells were treated with AT-00-11A-MMAE (5 nM), AT2604 (5 nM), or a combination of AT-00-11A-MMAE or AT2604 and KRASi (MRTX1133; 100 nM for AsPC-1, 10 nM for SW1990, 1 µM for PANC-1). Each data point represents a technical replicate; error bars denote mean ± s.d. Data is representative of 3 biological replicates. Statistical significance was determined by performing Welch’s t-test. **D.** Tumor volume growth curve of HPAC tumor-bearing mice treated with vehicle, AT-00-11A-MMAE or two different doses of AT2604 (n=6 mice per arm) twice weekly by IV injection. Statistical significance was determined by one-way ANOVA followed by Tukey’s multiple comparisons test. **E.** Immunoblotting analysis of ALPP/ALPPL2, phospho-ERK (Thr202/Tyr205), and ERK protein levels in PDAC PDOs treated with RASi (RMC-6236, 100 nM) for 48 hours. **F.** Top, relative viability of PDAC PDOs treated with a dose response of RMC-6236 alone or in combination with increasing doses of AT2604 for 6 days. Each data point represents the mean of triplicates ± s.d.; viability curves shown are representative of three independent experiments. Bottom, Highest Single Agent synergy score values and heatmaps from data represented above. Tiles in heat map represent HSA synergy scores (red = synergy, white = additivity, green = antagonism). **G.** IHC images of SW1990 and AsPC-1 tumors stained for ALPP/ALPPL2 or pERK (T202/Y204) treated with 10 mg/kg once daily RMC-6236 for 3 and 7 days. Scale bars = 50 µm. **H.** Quantification of human ALPP/ALPPL2 from serum of mice from **G** treated with 10 mg/kg once daily RMC-6236 for 3 or 7 days (RMC-6236) as measured by ELISA, circles = 3 days, triangles = 7 days. Significance was determined by Welch’s t test. Statistical significance was determined by performing Welch’s t-test. **I.** Average tumor volume of subcutaneous SW1990 xenograft tumors treated with AT-00-11A-MMAE (3 mg/kg IV QW x 2) and vehicle, AT2604 (3 mg/kg IV QW x 2) and vehicle, RMC-6236 (10 mg/kg PO QD x 14) and AT-00-11A-MMAE, or AT2604 and RMC-6236. n = 9-11 mice per arm. Each point represents mean tumor volume ± s.e.m. Statistical significance was determined using two-way ANOVA followed by Tukey’s multiple comparisons test. **J.** Percent change of individual tumor volumes on day 27 relative to day 0. Statistical significance was determined by one-way ANOVA followed by Tukey’s multiple comparisons test. **K.** Kaplan-Meier curve quantifying the probability of survival over time for each treatment arm. Statistical analysis was determined by Mantel-Cox test. *P<0.05, **P<0.01, ***P<0.001, ****P<0.0001; ns, not significant.

We next sought to determine if the observed internalization of AT2604, as a single agent and in combination with KRASi, resulted in an anti-proliferative phenotype in our panel of PDAC cell lines. First, we determined that our cell lines tested were sensitive to single agent, unconjugated MMAE (Supplementary Fig. S8B). Next, we treated PDAC cells with isotype control ADC (AT-00-11A-MMAE) or AT2604 alone or in combination with KRASi and found that the combination of AT2604 and KRASi had a significantly greater anti-proliferative effect than AT-00-11A-MMAE and KRASi (Fig. 6C). We also evaluated the single-agent efficacy of AT2604 in a high baseline ALPP/ALPPL2 expressing HPAC PDAC xenograft model demonstrating significant tumor stasis/regression, highlighting the efficacy of this ADC in vivo in the context of PDAC (Fig. 6D). We next sought to move into more advanced, clinically relevant models of PDAC, as well as using RMC-6236, the RAS(ON) multi-selective inhibitor with completed phase III clinical trials and FDA approval for second-line therapy in patients with advanced PDAC (9) (NCT06625320). Patient-derived organoids (PDOs) have emerged as a highly translational model that has been shown to more accurately recapitulate patient responses to therapies (51). We first aimed to determine if RAS inhibitor (RASi) treatment could increase ALPP/ALPPL2 expression in PDOs. We found that 2-day RASi treatment increased total ALPP/ALPPL2 protein levels (Fig. 6E) and *ALPP/ALPPL2* mRNA levels in PDAC PDOs (Supplementary Fig. S8C). Next, we found that this RASi-induced ALPP/ALPPL2 expression conferred a synergistic anti-proliferative phenotype when PDAC PDOs were treated with a combination of RASi and AT2604, but not RASi and the isotype control ADC (Fig. 6F, Supplementary Fig. S8D).

We next sought to determine if the observed ALPP/ALPPL2 increased expression following RASi was recapitualed *in* vivo. To this end, we treated mice harboring SW1990 or AsPC-1 CDXs with 10 mg/kg RMC-6236 once daily for 3 or 7 days. Upon performing IHC staining on these treated tumors, we observed a significant increase in ALPP/ALPPL2 expression that coincided with a loss of phospho-ERK staining at both timepoints (Fig. 6G, Supplementary Fig. S8E). We also collected serum from these mice and found that there was a significant increase in secreted ALPP/ALPPL2 following RASi treatment relative to vehicle (Fig. 6H), suggesting a potential serum biomarker to determine whether treatment with an anti-ALPP/ALPPL2 ADC may provide benefit (25). Finally, to determine if the enhanced efficacy of AT2604 provided by RMC-6236 could be extended to *in vivo* models of PDAC, we generated CDX tumors from SW1990 cells in nude mice and treated mice with AT-00-11A-MMAE, AT2604, AT-00-11A-MMAE with RMC-6236, or AT2604 with RMC-6236. All mice received two doses of 3 mg/kg ADC (AT2604 or AT-00-11A-MMAE) 7 days apart, alone or in combination with 10 mg/kg RMC-6236 once a day for two weeks. We found that the combination of AT2604 and RMC-6236 markedly delayed tumor outgrowth relative to either single agent treatment (Fig. 6I, Supplementary Fig. S8F), with no significant change in body weight observed (Supplementary Fig. S8G). At day 27, we found that the combination of AT2604 and RMC-6236 demonstrated significant tumor regression relative to all other treatment arms (Fig. 6J). Finally, we found that the combination treatment significantly improved the probability of survival relative to either treatment alone (Fig. 6K). Taken together, we conclude that concomitant AT2604 and RMC-6236 presents a potential therapeutic strategy for PDAC patients.

## DISCUSSION

Recently developed mutant-selective KRAS and RAS(ON) inhibitors have demonstrated remarkable responses for patients with advanced PDAC; however, as monotherapies, all these drugs are all limited by intrinsic or acquired resistance. Approximately 60% of acquired resistance to KRAS/RAS inhibitors can be attributed to genomic alterations, while the other ∼40% is thought to occur through nongenomic mechanisms (13,31). Here, we performed global proteome, phosphoproteome, and surfaceome analyses after acute and long-term KRASi treatment in PDAC models to identify nongenomic adaptations and associated vulnerabilities for combination therapy to prevent or delay therapeutic resistance and improve PDAC patient outcomes. Proteomics datasets generated in this study identified essential proteins and pathways downregulated at 2 days of KRASi treatment, as previously observed (2,3); followed by a rebound at 7 days of sustained KRASi treatment with a return to baseline in PDAC models treated to resistance, consistent with our previous findings demonstrating early loss of KRAS-dependent signaling followed by compensatory pathway reactivation as cells acquire drug tolerance (14). These proteomic datasets align with our prior efforts at modeling proteomic adaptations to KRAS^G12C^ inhibition with pre-clinical inhibitors in that much of the acute adaptations can be characterized as a cytostatic response and survival program with associated downregulation of cell cycle, transcription, and translation concurrent with activation of compensatory stress and metabolic pathways, most prominently an early and sustained upregulation of lysosomal metabolism (14). The temporal evolution of KRASi adaptive signaling pathways as determined by our phosphoproteomic analyses demonstrate that PDAC cells survive the initial KRASi-induced proliferative block by engaging pro-survival, motility, and EMT-associated signaling. In sum, these proteomic and phosphoproteomic datasets provide a temporal roadmap to nongenomic adaptations during development of KRASi resistance and a resource for identification of targetable pathways to prevent adaptation and drive deeper response to KRASi.

Using surface-enriched proteomics, we further characterized how the KRASi-induced surfaceome changes temporally, with the goal of finding an upregulated protein for ADC targeting to enhance the efficacy of KRASi. We detected upregulated surface targets that are currently undergoing clinical evaluation with ADC targeting, such as MUC1 (52) (NCT05875168), ITGB6 (NCT04389632), and CEACAM5 (NCT06710132). Combining our experimental dataset with published expression databases through a machine learning algorithm, we nominated and validated several PDAC surface proteins that are upregulated after KRASi-treatment and possess characteristics of an actionable ADC target, such as low normal tissue expression and high tumor expression. Through this analysis, ALPP/ALPPL2 emerged as a top candidate due to the significant induction of surface expression and limited healthy tissue expression. Mechanistically, RAS-MAPK inhibition leads to increased active phospho-STAT3 (44), a transcription factor that binds the promoter region of the ALPP and ALPPL2 genes in non-small cell lung cancer (43). We found that genetic ablation of STAT3 or pharmacological inhibition of STAT3 activity by JAK inhibitor treatment could prevent the upregulation of ALPP surface expression following KRASi. Recent studies have also implicated activation of FoxO3a as a transcriptional regulator of ALPP upregulation in response to EGFR inhibition in lung adenocarcinoma, suggesting the mechanism of upregulation may be context-dependent (27). Beyond these cell-intrinsic mechanisms, the PDAC tumor microenvironment is a rich source of paracrine STAT3-activating cytokines, facilitated by cancer-associated fibroblasts that secrete IL-6, which drive STAT3 activation and disease progression in PDAC models (53). Given that our cell line and organoid models lack this stromal compartment, future work should aim to determine whether the tumor microenvironment contributes to, or potentiates, KRASi-induced ALPP/ALPPL2 upregulation.

We and others have developed anti-ALPP/ALPPL2 ADCs for a variety of cancer types (22,26,27). AT2604 represents a potential best-in-class ALPP/ALPPL2 targeting ADC with next-generation engineering including homogenous drug antibody ratio (DAR 4) and site-specific MMAE conjugation. These features translate into a potent and specific anti-ALPP/ALPPL2-MMAE ADC with single-agent efficacy in *in vivo* models of gastric, gastroesophageal, and PDAC cancers. Ultimately, we sought to use AT2604 to therapeutically exploit the KRASi-induced upregulation of surface ALPP/ALPPL2 expression. Indeed, we found that the combination of KRASi/RASi and AT2604 significantly impaired the growth and viability of PDAC cell lines and patient-derived organoids. This favorable anti-proliferative phenotype was extended to *in vivo* models of PDAC in which we demonstrated that the combination treatment significantly delayed tumor outgrowth relative to single-agent treatments. Notably, our work extends beyond PDAC with ALPP/ALPPL2 upregulation in response to KRAS/MAPK pathway inhibition as a conserved feature in KRAS-mutant (and EGFR-mutant) cancers, including ovarian and lung cancer (companion study), thus suggesting a KRAS-mutant and MAPK-pathway driven tumor agnostic applicability of our work. Independent work similarly identified EGFR inhibition-induced ALPP/ALPPL2 upregulation and combination efficacy of EGFR inhibition and a pre-clinical ALPP/ALPPL2-targeting ADC (27).

Our work highlights an important temporal consideration for the preclinical and clinical evaluation of KRASi and ADC combination therapy. As ALPP/ALPPL2 were most upregulated at the 2 day and 7 day timepoints (Fig. 3A, 4A), we observed a significant response in both cell, organoid, and mouse models when administering KRASi and AT2604 simultaneously. This concomitant approach provided the advantage of exploiting the more uniform acute adaptation to RASi at 2 days compared to the divergent phenotypes at resistance (Fig. 1B, 1C, Fig. 3A, Supplementary Fig. S5A), where upregulation of cell surface targets may be more predictable in early response to KRASi in patients. Furthermore, as KRASi treatment has been shown to increase the sensitivity of cells to cytotoxic agents (31) and their lysosomal metabolism (Supplementary Fig. S2D), further investigation into how different ADC payloads and linkers can exploit the vulnerable cellular state of acute RASi adaptation may also prove fruitful for KRASi and ADC combination therapies. Future work identifying cell surface targets associated with KRASi resistance or prolonged exposure to KRASi will be crucial for expanding the applicability of KRASi and ADC combination therapies to PDAC patients who have already developed resistance to KRASi treatment.

One limitation of this combination approach is that PDAC cell lines with little to no baseline ALPP/ALPPL2 expression had no appreciable upregulation of ALPP levels, suggesting the necessity of at least some detectable baseline ALPP expression for KRASi-induced upregulation. This is similar to recent work identifying HER2 as an upregulated cell surface target in response to pan-RAS(ON) inhibition in select PDAC models with robust combination activity in models with higher baseline HER2 but no combination activity in models with low HER2 levels basally (13,54). Future work aims to nominate other surface targets from our datasets to target low-ALPP/ALPPL2 PDAC tumors such as MUC1 targeting by DS-3939a (52) or an alternative Claudin 18.2 ADC (55). Alternatively, bi-specific antibody-targeting strategies to harness other KRASi-induced targets (e.g. MUC1) may mitigate low/absent ALPP/ALPPL2 levels for more effective targeting of inter- and intra-tumoral surface expression heterogeneity. Additional considerations include selecting different cytotoxic payloads for conjugation to our anti-ALPP antibody, CAR-T approaches for an orthogonal approach for low-ALPP-expressing cancer cells, and determining if the transcriptional subtypes of PDAC demonstrate differential sensitivities to these treatments. While ADCs have not worked to date in PDAC clinically, such as HER2 targeting (56) and Claudin 18.2 targeting (57) (NCT03816163), this may be attributed to barriers to ADC delivery (58) as opposed to target selection. As KRASi targeting demonstrates significant stromal remodeling (59,60), future work will determine whether this improves ADC delivery and thereby the potency of this approach. Altogether, our work provides a crucial proof of concept for KRASi and ADC combination therapy design and highlights AT2604 anti-ALPP/ALPPL2 ADC as one such promising therapeutic strategy to drive deeper, more sustained anti-tumor responses and potentially forestall KRASi resistance. Furthermore, our work provides a detailed roadmap for designing rational KRASi and ADC combination therapies, from target identification to therapeutic design and preclinical evaluation.

## METHODS

### Cell Culture

AsPC-1, Panc1, Panc0203, Panc0403, HPAC, and SW1990 cells were obtained from ATCC, and SUIT2, KP-4, KP-1N were obtained from the JCRB Cell Bank. PK45H and PK-1 cells were obtained from the RIKEN BRC. Cell lines were maintained in a centralized cell bank, authenticated by assessment of cell morphology as well as short tandem repeat fingerprinting, and routinely inspected for Mycoplasma contamination using PCR. After thawing, cell lines were cultured for no longer than 30 days. Cell lines were maintained at 37°C with 5% CO2 and grown in DMEM (Panc1, HPAC, and SW1990) or RPMI 1640 (AsPC-1, Panc0203, Panc0403, SUIT2, PK45H, KP4, PK-1, and KP-1N) supplemented with 10% FBS and 1% penicillin/streptomycin, unless otherwise specified.

### Generation of Isogenic Models of Acquired Resistance to MRTX1133

MRTX1133-resistant cell lines were generated as previously described (31). Briefly, PDAC lines were treated with a continuous dose escalation from as low as 1 nM to 10 µM. Once resistance to MRTX1133 was confirmed, resistant cell lines were routinely maintained in the corresponding medium with 1 µM (10% of the maximum dose reached during the evolution of resistance) of MRTX1133.

### Surfaceome enrichment

Surfaceome enrichment was performed as previously described (34) with some modifications. Cells were seeded in 15 cm dishes for at least 48 h before labeling, and corresponding treatments were applied, aiming for 50 million cells at 90% confluency at the day of proteomics experiments. Culture dishes were washed once with 10 mL cold PBS, then incubated with 5 mL 1mM EDTA in PBS for approximately 5-15 min depending on the cell line. Cell suspension was collected and washed 2x with 10 mL cold PBS by spinning at 1000 x g at 4°C for 2 min. Cell pellets were resuspended in labeling mix containing 200 μM aminooxy-biotin, aniline (1:10) and 1 mM sodium periodate in PBS pH 6.5. The cell suspension was incubated at 4°C rotating at 10 RPM for 30 min. The labeling mix was removed by spinning and pellets were washed 2x with 15 mL PBS pH 7.4 to remove excess labeling solution. Washed cell pellets were lysed in 500 μL NP-40 lysis buffer (50 mM Tris HCl, pH 8.0, 150 mM NaCl, 1% IGEPAL CA-630) plus Halt Protease Inhibitor by rotating at 4°C at 30 RPM for 1 hour and clarified at 20,000 x g at 4°C for 15 min. 100 µL streptavidin agarose beads (200 µL slurry) were washed 2x with PBS and added into each lysate, then incubated at 4°C rotating at 10 RPM for 16 h. The next day, agarose beads were pelleted and washed 2x with 4 M Urea. Then beads were incubated with EPPS containing 10 mM TCEP for 30 min, followed by addition of 10 mM IAA and incubation for 30 min at room temperature. After reduction and alkylation, beads were washed 3x with MS-grade water. 1 µg trypsin and 1 µg LysC in 50 µL 200 mM EPPS (pH 8.0) were added into each sample, and the samples were incubated at 37°C mixing at 1200 RPM for 16 h. The next day, 3 µL TMTpro stock (20 mg/ml) and 15 µL acetonitrile (ACN) were added into each sample, then incubated at 21°C mixing at 1200 RPM for 60 min. TMTpro labeled samples were quenched via 7 µL hydroxylamine stock (5%) at 21°C mixing at 1200 RPM for 10 min. Quenched samples were pooled together. 10% formic acid (FA) was added to acidify TMTpro-labeled samples and was desalted via C18 Sep-Pak (60 mg). Desalted peptides were eluted from Sep-Pak with 70% ACN and 1% FA and dried via SpeedVac. Dried peptides were dissolved in 10 µL 5% can and 5% FA. 10 µL peptides were analyzed in an Thermo Scientific Orbitrap Exploris 480 coupled with a nano-LC separated in a 120 min gradient. MS acquisition was performed in data-dependent (DDA) high-resolution MS2 mode and peptide ions were pre-selected via FAIMS Pro Duo before the MS inlet.

### Whole cell proteomics and phosphoproteomics

Cell pellets were lysed in 8 M urea in 200 mM EPPS pH 8.0 plus Halt Protease and Phosphotease Inhibitor Cocktail. Protein concentration of each lysate was determined via BCA, then 50 µg protein was aliquot from each sample and normalized to 1 µg/µL using lysis buffer. Protein reducing and alkylation were performed with 10 mM TCEP and 20 mM IAA by incubating at 37°C for 45 min. Protein was cleaned via the SP3 method: 10 µL SP3 beads and 60 µL ethanol were added into each sample and rotated at 21°C at 10 RPM for 10 min, then SP3 beads were washed 2x with 80% v/v ethanol. 1 µg trypsin and 1 µg LysC in 50 µL EPPS (200 mM pH 8.0) were added into each sample and was incubated at 37°C mixing at 1200 RPM for 16 h. The next day, 5 µL TMTpro stock (20 mg/mL) and 15 µL ACN were added into each digest and incubated at 21°C mixing at 1200 RPM for 60 min. The TMTpro reaction was quenched via 0.5% hydroxylamine at 21°C mixing at 1200 RPM for 10 min. 2 µL labeled samples were pooled in 200 µL 1% FA, then desalted via C18 Stage-Tip. Cleaned peptides were injected for a 45 min gradient DDA hrMS2 acquisition and the data was used to determine the ratio of peptide quantity in each TMTpro channel. Equal amount TMTpro-labeled peptides were pooled together and desalted via C18 Sep-Pak (60 mg). Peptides were eluted with 70% ACN and 1% FA and dried via SpeedVac. Phophopeptides were isolated from dried peptides according to the manufacturer’s protocol using a Thermo Fe-NTA Phosphopeptide Enrichment Kit (ThermoFisher, A32992). Enriched phosphopeptides were desalted via C18 Stage-Tip and dried in a SpeedVac. Phosphopeptides were injected into a Thermo Scientific Orbitrap Eclipse Tribrid coupled with nano-LC at 120 min gradient. The MS Instrument acquired in DDA hrMS2 mode with FAIMS Pro Duo. After phosphopeptide enrichment, the flow-through was dried in a SpeedVac and fractionated offline into 24 fractions through basic pH reverse phase C18 column. These 24 fractions were desalted via C18 Stage Tips and 12 fractions were analyzed in a Thermo Scientific Orbitrap Exploris 480 coupled with a nano-LC separated in a 90 min gradient. MS acquisition was performed in DDA high-resolution MS2 mode and peptide ions were pre-selected via FAIMS Pro Duo before the MS inlet.

### Proteomics Data Processing

All raw mass spectrometry files were jointly searched using the MaxQuant (61) (version 2.6.7.0) Andromeda search engine (62) and the UniProt Human Reference Proteome with known contaminants included (63) (20,417 sequences, accessed January 2025). The peptide length was set to 8-25 with a maximum mass of 4,600 Da. False Discovery Rate (FDR) for peptide identification was set at <0.01 with a minimum of one razor peptide. MS2 reporter ion was searched using manufacturer’s isotope correction factors for TMTpro-18plex (Thermo) with a reporter mass tolerance of 0.003 Da. For modified peptides, reporter ions were filtered by a minimum precursor intensity fraction (PIF) of 0.75, variable modifications were set to oxidation, N-terminal acetylation, and phospho–serine-threoninetyrosine (STY), and carbamidomethyl was set as fixed modifications with a maximum number of five modifications per peptide. Digestion was set to Trypsin/P with three maximum missed cleavages. Default Orbitrap settings were used for the spectrometer. For unmodified peptides, oxidation and N-terminal acetylation were set as variable modifications and carbamidomethyl was set as a fixed modification.

### Normalization and Differential Expression Analysis

The output files from MQ were imported into R (version 4.5.3) for all downstream data processing, filtering, and statistical analyses. For phosphoproteomic datasets, sites were filtered to retain high-confidence Class I phosphosites (localization probability > 0.75). Contaminants and reverse sequences were removed, and reporter ion intensities were sum-normalized within each cell line and individual TMT multiplex set. Differential expression analysis was performed using the limma (64) (3.66.0) package in R. To identify conserved proteomic and phosphoproteomic alterations across the diverse panel of models, a blocking term for each cell line was used to correct for baseline intensities (65). Missing data was not imputed. Principal component analysis (PCA) was performed using the factoextra (66) package in R, utilizing zero-filling for missing values and mean-centering. Pairwise sample similarity was evaluated using Pearson’s correlation coefficients, and the resulting correlation matrices were subjected to unsupervised hierarchical clustering applying the McQuitty linkage method (67).

### Database Integration and Metadata Annotation

Genetic dependencies were evaluated using the Cancer Dependency Map (DepMap). DepMap datasets and mutation calls (22Q2) were accessed via the AnnotationHub R package (68). Cell lines were filtered to isolate KRAS-mutant pancreatic ductal adenocarcinoma (PDAC) models, and median dependency scores were calculated for each gene across KRAS-mutant PDAC cell lines. Subcellular localization was annotated using the COMPARTMENTS database (69). Proteins were mapped to major cellular compartments (Nucleus, Cytoplasm, Mitochondrion, Lysosome, Plasma membrane, Cytoskeleton, Endosome, Peroxisome, and Golgi apparatus) using the maximum localization score for each gene, allowing for multi-compartment associations. Functionally validated phosphosites were annotated using the Regulatory Sites dataset from PhosphoSitePlus (32). To define the cell surfaceome, we compiled a consensus list of surface-localized proteins by integrating annotations from three comprehensive surfaceome datasets: the Cancer Surfaceome Atlas (36), the *in silico* human surfaceome (35), and PDAC surfaceome profiling (37). Finally, to prioritize targets for therapeutic translation, proteins with clinically investigated antibody-drug conjugates (ADCs) were manually curated and annotated using recent comprehensive clinical reviews (16).

### Gene Set Enrichment Analysis (GSEA) and Pathway Trajectories

Pathway enrichment was performed using the fgsea (70) and msigdbr (71) R packages. To account for both the magnitude and significance of protein regulation, proteins were ranked for GSEA using a composite metric: the log_2_(fold-change) multiplied by the min-max scaled −log_10_(adjusted *p*-value). Gene sets evaluated included canonical MSigDB libraries in addition to empirically defined KRAS-ERK (2,3) and MYC (29) specific signatures. Proteins were further mapped to established classical and basal pancreatic cancer transcriptional subtypes using defining gene sets previously described (72).

To model the temporal evolution of these pathways, GSEA net enrichment scores (NES) across all timepoints were clustered using Ward’s minimum variance hierarchical clustering. The optimal number of temporal clusters was determined empirically using the gap statistic method in the cluster R package (73). Pathway similarities were determined by computing the Szymkiewicz-Simpson similarity index using the leading-edge genes derived from the GSEA results for each pathway, followed by consolidation via Ward’s hierarchical clustering.

### Kinase Motif Profiling and Kinase-Substrate Networks

To deduce dysregulated kinase signaling, differentially regulated phosphosites were evaluated for kinase motif enrichment utilizing the Kinase Library (33). Relative kinase activities were quantified by calculating the product of the min-max normalized adjusted *p*-value and the log_2_ enrichment factor for the most significant enrichment direction of each respective kinase. Dynamic kinase-substrate networks were then constructed using a previously established methodology (3). Briefly, a composite kinase-substrate interaction score was calculated for each phosphoprotein by summing multiple weighted lines of evidence: (i) a motif-based score derived from the Kinase Library (integrating phosphosite motif preference with the dataset-specific altered kinase activity measure), (ii) previously reported, empirically validated kinase-phosphosite relationships from PhosphoSitePlus (74) , and (iii) documented protein-protein interactions sourced from PhosphoNetworks and BioGRID (v4.4.213) (75). To visualize the most robust signaling nodes, constructed networks were filtered to retain only interactions with an absolute composite score greater than one standard deviation. Finally, network nodes were annotated with their corresponding median genetic dependency scores derived from the DepMap KRAS-mutant PDAC subset, as described above.

### Functional Overrepresentation Analysis

Functional categorization of the dysregulated phosphoproteome was evaluated through overrepresentation analysis of signaling pathways and subcellular compartment localizations. Statistical enrichment for each category was determined using a one-tailed hypergeometric test, requiring a minimum background set of five phosphoproteins per tested category. The resulting *p*-values were corrected for multiple hypothesis testing using the Benjamini-Hochberg method.

### Integration of Surface-Enriched and Global Proteomics

BAMX surface-enriched proteomics and global proteomics were integrated using Stouffer’s weighted z-score method. For each protein, dataset-specific z-scores were weighted by the square root of their respective sample sizes and summed. The resulting statistic was normalized by the square root of the sum of the squared weights, yielding a combined z-score following a standard normal distribution under the null hypothesis. Combined z-scores were converted to two-sided *p*-values using the standard normal cumulative distribution function, with effect directions preserved by signing individual z-scores prior to summation. This weighting scheme provided proportionally greater power to statistically powered datasets while maintaining a closed-form null distribution for hypothesis testing.

### Multi-Omic Feature Engineering

To systematically prioritize ADC targets, a 10-feature matrix was constructed encompassing healthy tissue profiles, cancer expression, and experimental KRASi dynamics (Supplemental Table S8). The median normal-tissue RNA expression (log_2_(TPM + 1)) was obtained from the Human Protein Atlas (HPA) consensus dataset (38), excluding placental tissue to reflect the adult oncology patient population. HPA immunohistochemistry (IHC) data were utilized to calculate normal and tumor protein expression features: the normal IHC score was defined as the proportion of profiled healthy tissues with appreciable (medium/high) staining, while the pancreatic cancer IHC score was defined as the fraction of tumors with any detectable staining. Pancreatic ductal adenocarcinoma prognostic significance from the HPA Kaplan-Meier analyses (39) was converted to binary (1 for log-rank *P* < 0.0001; 0 otherwise). Tumor RNA expression was derived from the TCGA-PAAD cohort as the median log_2_(TPM + 1) across all tumors, and tumor-versus-normal protein log_2_(fold-change) was extracted from the CPTAC pancreatic ductal adenocarcinoma dataset (76).

Experimental features capturing KRASi-induced molecular dynamics included: (i) acute 24-hour MRTX1133 treatment RNA-seq log_2_(fold-change) in AsPC-1 cells (40), (ii) the median global proteomic log_2_(fold-change) across 2-day and 7-day MRTX1133 treatments, and (iii) the median BAMX-enriched surfaceome log_2_(fold-change) across 2- day and 7-day MRTX1133 treatments. Summarizing the proteomic temporal data by median fold-change reduced model dimensionality and prioritized robust, reproducible target induction. Finally, genes were categorically annotated for cell surface localization and known clinical ADC status as previously described.

### Ridge Logistic Regression Modeling and Target Scoring

A ridge-penalized logistic regression model was employed to calculate a continuous ADC target-prioritization score. A positive training set was defined by filtering known, clinically investigated ADC targets for those demonstrating positive KRASi-mediated induction (log_2_(fold-change) > 0.25) in at least one experimental modality (RNA-seq, global proteomics, or surfaceomics). The 31 targets meeting these criteria were labeled as positive training examples (Class = 1). All remaining genes were labeled as 0; notably, this negative class represents the absence of current clinical ADC development rather than confirmed biological unsuitability. Ridge regularization (glmnet package in R, a = 0) was explicitly selected to shrink correlated features toward zero thereby minimizing overfitting. The 10-feature matrix, excluding known ADC status and training ID, for genes with complete cases was standardized to zero mean and unit variance. The optimal regularization parameter (l) was selected via five-fold cross-validation using binomial deviance as the optimization metric, utilizing the conservative l_1se_ value.

To score the full candidate gene set, feature vectors were standardized using the centering and scaling parameters derived from the complete-case training matrix, with any missing values zero-filled. The model-generated linear predictor (LP), representing the log-odds score, was calculated for each gene. To provide an intuitive, relative prioritization metric, scores were min-max normalized and transformed to a [−1, +1] scale:

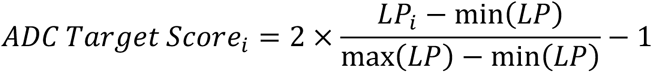

where *_LPi_* is the ridge-model linear predictor for gene *i*.

### Mouse Models

Animal experiments were performed in accordance with protocol 10-055 approved by the Institutional Animal Care & Use Committee (IACUC) at Dana-Farber Cancer Institute (DFCI). Athymic nude female mice (NU/J, Homozygous) were obtained from the Jackson Labs at 7-weeks old (Stock # 002019).

For the cell-derived xenograft (CDX) studies in Figures 6G-K and Supplementary Figures S8E-G, subcutaneous tumor implantation was performed as follows: Cells were harvested by Trypsin and neutralized upon detaching with media supplemented with 20% FBS. Cells were pooled from multiple plates and centrifuged at 800 x g for 3 minutes. Supernatant was aspirated and resuspended in 10 mL of complete media. Cells were counted, and viability was monitored to ensure cell health before tumor implantation. The required number of cells was moved to a new tube and pelleted. Upon aspirating the supernatant, the cell pellet was resuspended in ice-cold PBS, and then Matrigel (Corning, 356231) was added at a 1:1 ratio to a final concentration of 5 million cells per 100 µL. Mice were injected subcutaneously with 5 million cells. Tumors were allowed to form for 2-3 weeks until tumor volume reached between 200 – 300 mm^3^, at which point mice were randomized into indicated treatment groups. For the SW1990 CDX tumor growth inhibition (TGI) study in Figures 6G-K and Supplementary Figures S8E-G, mice were treated with vehicle (AT-00-11A, 3 mg/kg IV QW x 2 and DMSO/PEG400/HS15/Water PO QD x 14), AT2604 (3 mg/kg IV QW x 2 and DMSO/PEG400/HS15/Water PO QD x 14), RMC-6236 (MedChemExpress, 10 mg/kg PO QD x 14 in DMSO/PEG400/HS15/Water and AT-00-11A, 3 mg/kg IV QW x 2), or a combination of AT2604 and RMC-6236. Tumor volumes were monitored by caliper measurement every 3 – 4 days during and after treatment until humane endpoint was reached. For the AsPC-1 and SW1990 CDX pharmacodynamic experiments in Figure 6G-H and Supplementary Figure S8E, mice were treated with vehicle (DMSO/PEG400/HS15/Water) or 10 mg/kg RMC-6236 PO QD for either 3 or 7 days. Tumors were collected 4 hours after the final dose and fixed in 10% formalin solution (Sigma Aldrich, HT501320) for 48 hours, washed twice with PBS, and stored in 70% ethanol at 4°C.

The following procedures were approved by the Institutional Animal Care and Use Committee (IACUC) of WuXi AppTec (Nantong) Co., Ltd. and conducted in accordance with Association for Assessment and Accreditation of Laboratory Animal Care (AAALAC) guidelines. Animal care complied with institutional guidelines and AAALAC recommendations; animals were observed at least once daily for clinical signs, and humane endpoints were applied as described below.

The human gastric carcinoma (NCI-N87) and human pancreatic ductal adenocarcinoma (HPAC) were used for in vivo efficacy evaluation in Figures 5E and 6D, respectively. Cells were maintained in appropriate medium supplemented with 10% fetal bovine serum (Dong Ling) and 1% Antibiotic-Antimycotic (Gibco) at 37°C in a humidified incubator with 5% CO_2_. Cells were sub-cultured twice weekly, and exponentially growing cells were harvested using 0.25% trypsin-EDTA (Gibco) and counted prior to tumor inoculation. Female BALB/c Nude mice (*Mus musculus*), 6–8 weeks old and weighing 18–22 g, were obtained from Shanghai Lingchang Biotechnology Co., Ltd. (Shanghai, China). Animals were housed in individually ventilated polycarbonate cages (375 mm × 215 mm × 180 mm) with corn cob bedding (changed twice weekly), with six mice per cage, under controlled environmental conditions (temperature 20–26°C, relative humidity 40–70%) and a 12-hour light/dark cycle. Mice had ad libitum access to irradiation-sterilized pelleted chow and sterile drinking water. For the NCI-N87 CDX efficacy study in Figure 5E, n = 6 mice per arm were treated with vehicle, 3 mpk isotype ADC, or indicated concentrations of AT2604 intravenously for 2 doses once a week. For the HPAC CDX efficacy study in Figure 6D, n = 6 mice per arm were treated with either vehicle, AT-00-11A-MMAE, or indicated concentrations of AT2604 for 2 doses once a week. Tumors were measured by caliper every 3-4 days for both studies as well as body weight, and were continued until humane endpoint was reached.

For PDX studies detailed in Figures 5F-G and Supplementary Figures S7D-E, two different subcutaneous HuPrime® xenograft cancer models were used in female BALB/c Nude mice: a human gastric carcinoma, GA2419, and a human gastroesophageal junction cancer (GEJ), GA6203. For both PDX efficacy studies, three mice per arm were treated with either vehicle or indicated concentration of AT2604 for 3 doses once a week. Tumor volumes were measured every 3-4 days by caliper measurements along with body weight until humane endpoint was reached.

Female BALB/c nude mice (*Mus musculus*, BALB/c nude, 6–8 weeks old at inoculation) were obtained from Jiangsu GemPharmatech Co., Ltd. (Jiangsu, China). Upon arrival, animals were acclimatized before study initiation and housed at the Crown

Bioscience Beijing/Taicang facilities accredited by AAALAC. Mice were maintained in individually ventilated polysulfone cages (approximately 325 × 210 × 180 mm) with autoclaved crushed corncob bedding, changed weekly, with up to five mice per cage. Environmental conditions were controlled at 20–26 °C, 40–70% relative humidity, and a 12 hour light/dark cycle. Standard irradiated rodent chow and autoclaved, filtered reverse-osmosis water were provided ad libitum. Environmental enrichment (tissue paper, cardboard cylinders, chew sticks, plastic clubhouse toys) was provided for all cages. All in vivo procedures were reviewed and approved by the Institutional Animal Care and Use Committee (IACUC) of Crown Bioscience prior to study initiation.

### Therapeutic Sensitivity Profiling of Cell Lines

Cells were seeded into a 96-well plate (500 - 1,000 cells/well, Corning; Cat. # 3903) in 50 µL complete medium on Day 0. Cell lines were allowed to seed overnight and then treated with indicated doses of MRTX1133 (acquired from Mirati/Bristol Myers Squibb), RMC-6236 (MedChemExpress), MMAE, or DMSO (Day 1). Cell viability was assessed on matched untreated “Day 0” plates to obtain a baseline cell viability. Treated plates were incubated for 5 days on treatment, and then cell viability was assessed using a Cell-TiterGlo 2.0 Cell Viability assay (Promega; Cat. # G9241), executed according to the manufacturer’s instructions. Fluorescence was read using a CLARIOstar microplate reader. Technical triplicates were conducted for each biological replicate, and a total of at least three biological replicates were done for each cell line. The GR_50_ value was calculated for all 2-D cell line experiments using the following equation:

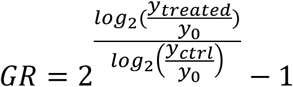

where y_0_ = Day 0 signal, y_ctrl_ = Control (0 nM) mean signal at Day 5, y_treated_ = Treated mean signal at endpoint as previously described (77).

### Western Blotting

Cells were washed twice on ice with PBS, scraped and collected in prechilled Eppendorf Tubes. Cell pellets were lysed in RIPA buffer (50 mM TrisHCl pH 7.4, 150 mM NaCl, 2 mM EDTA, 1% NP-40, 0.1% SDS) supplemented with protease and phosphatase inhibitor cocktails (Thermo Scientific; 78446) and 1 mM DTT. Lysates were clarified by centrifugation at 14,000 g for 15 minutes at 4°C followed by protein concentration quantification by the Micro BCA™ Protein Assay Kit (Thermo Scientific; 23235). Equal amounts of protein were separated by SDS-PAGE electrophoresis and transferred to nitrocellulose membranes. Membranes were blocked in 5% (w/v) BSA in TBS with 0.1% Tween 20 (TBS-T) (Goldbio, A-420-100) for 1 hour and incubated overnight at 4°C in primary antibody. Membranes were washed three times with TBS-T and then incubated with peroxidase-conjugated secondary antibody diluted in 2% BSA in TBS-T for 1 hour at room temperature. Protein levels were detected using enhanced chemiluminescence (ECL) developing reagent (Thermo Fisher, 32106). The antibodies used are as follows: ALPP (Abcam, ab133602, 1:10000), p-ERK (T202/Y204, Cell Signaling Technology, 9101, 1:1000), ERK (Cell Signaling Technology, 9102, 1:1000), p-STAT3 (Y705, Cell Signaling Technology, 9145, 1:1000), STAT3 (Cell Signaling Technology, 4904, 1:1000), Actin (Sigma, A5441, 1:10000), vinculin (Millipore, 05-386, 1:5000), FLAG (Cell Signaling Technology, 14793, 1:1000), Mouse-HRP (Promega, W4021, 1:5000), and Rabbit-HRP (Thermo, 31460, 1:5000).

### Patient-derived organoids

All patients from whom PDAC organoid models were derived in this study provided written and informed consent as per IRB-approved protocols at Dana-Farber Cancer Institute, permitting access to their clinical and genomic data (DFCI 03189, 14-408). All organoids utilized in this study were annotated with genomic and clinical information and subsequently utilized in a de-identified, HIPAA-compliant manner as previously described (31). The human PDAC patient-derived organoids (PDOs) were cultured at 37°C in 5% CO_2_. PDOs were grown in growth factor-reduced Matrigel (Corning, 356231) domes and cultured with human complete OPAC media: Advanced DMEM/F12-based-conditioned medium, 1 x B27 supplement, 10 mM HEPES, 1.25 mg/mL Primocin, 100 µM Pen/Strep, 2 mM Glutamax, 10 mM nicotinamide, 1.25 mM n-Acetyl cysteine, 50 ng/mL EGF, 100 ng/mL hFGF10, 10 nM hGastrin I, 500 nM A83-01, 100 ng/mL Noggin, 1 x Wnt-3A conditioned 10% FBS DMEM (50% by volume), and 1 x R-spondin Conditioned Basal Medium (10% by volume).

For PDO drug testing, organoids were dissociated using TrypLE Express (Thermo Fisher, 12604021) supplemented with DNase at 1:200 and seeded (1,000 – 2000 cells/well) into 384-well black ultra-low attachment plates (Corning, 4588) in 20 µL OPAC media with 10% Matrigel and 1:100 ROCK inhibitor (Y-27632). Cells were allowed to seed overnight and then treated with the indicated drugs. Following 6 day drug incubation, cell viability was assessed using a CellTiter-Glo® 3D Viability assay (Promega, G9683) according to the manufacturer’s instructions. Luminescence was measured using a FLUOstar Omega microplate reader. Technical triplicates were conducted for each biological replicate, and a total of three biological replicates were done for each PDO line tested. PDO dose response data were analyzed using standard relative viability nonlinear regression. Luminescence values were normalized to vehicle treated controls and drug concentrations were log-transformed. Normalized viability data were fit by nonlinear regression (log[inhibitor] vs. response, variable slope, four parameters) in GraphPad Prism to generate dose response curves and determine IC50 values. Synergy was assessed using (78).

For PDO protein extraction, organoids were dissociated using TrypLE Express (Thermo Fisher, 12604021) supplemented with DNase at 1:200 and seeded (7.5 x 10^5^ cells/well) into 6-well ultra-low attachment plates (Corning, #CLS3471) in 2 mL of OPAC media supplemented with 10% Matrigel and 1:100 ROCKi. Cells were allowed to seed for 48 hours, and then 100 nM RMC-6236 or vehicle was added to PDOs for 48 hours. Following drug incubation, PDOs were collected and centrifuged at 1,500 rpm for 5 minutes at 4°C. The supernatant was aspirated, and the samples were washed twice with ice-cold PBS before resuspending in 10 mL of Cell Recovery Solution (Corning, 354253) and incubating on ice for 30-60 minutes. The samples were washed again with ice-cold to ensure Matrigel is fully removed, resuspended in 1 mL ice-cold PBS, transferred to a 1.5 mL Eppendorf tube, and spun down at 600 x g for 10 minutes at 4°C. All supernatant was aspirated and pellets were snap frozen and stored at −80°C. Protein extraction and immunoblotting were performed as described above.

### RNA Extraction

Total RNA was isolated from cell pellets using TRIzol Reagent following liqud-liquid extraction protocol. Briefly, TRIzol reagent was added to the cell pellet and homogenized by vortexing for 30 seconds and allowed to incubate for 5 minutes. Next, 200 µL of chloroform was added per 1 mL of TRIzol, vortexed for 30 seconds, incubated for 2-3 minutes, and then centrifuged at 12,000 x g for 15 minutes at 4 °C. The upper aqueous phase was transferred to a new tube. RNA was precipitated by adding 500 µL of isopropanol, vortexing for 30 seconds, and incubating at room temperature for 10 minutes. Samples were then centrifuged at 12,000 x g for 10 minutes at 4°C. The supernatant was discarded and the RNA pellet was resuspended in 1 mL of 75% ethanol per 1 mL TRIzol reagent used. Samples were vortexed and centrifuged at 7,500 x g for 5 min at 4°C. The supernatant was discarded and the RNA pellets were allowed to air dry for 20-30 minutes in a chemical fume hood to ensure complete ethanol evaporation. The purified RNA pellet was resuspended in 20-50 µL RNase-free water, pre-heated to 55-60°C, and mixed well by pipette. Complementary DNa (cDNA) was synthesized from 500 ng RNA per reaction using the High-Capacity cDNA Reverse Transcription Kit (Applied Biosystems, #4368814) following the manufacturer’s instructions. Briefly, reverse transcription was performed on a themocycler using the following conditions: 25°C for 10 min, 37°C for 120 min, 85°C for 5 min, and hold at 4°C. cDNA was diluted in nuclease-free water as needed to a working concentration of 60 ng per 9 µL. qPCR reactions were assembled in 20 µL aliquots containing 9 µL diluted cDNA (60 ng), 0.5 µL forward primer (Sequence: CAACGAGGTCATCTCCGTGATG), 0.5 µL reverse primer (Sequence: ACACGGTGAACCGCAACTGGTA), and 10 µL SYBR Green master mix and run on a QuantStudio 7 Flex System. Relative mRNA expression of ALPP/ALPPL2 were normalized to the housekeeping gene L32.

### Flow Cytometry

Cells were seeded in 6-well plates and treated with the indicated conditions. Following treatment, cells were washed with cold PBS and collected with TrypLE Express Enzyme (Gibco, 12604021). Cells were pelleted at 800 x g for 3 minutes and resuspended in cold FACS buffer (Thermo Fisher, 00-4222-26) (1% FBS in PBS). The cells were then washed twice with cold FACS buffer and incubated in primary antibody diluted in FACS buffer on ice in the dark for 30 minutes. If an unconjugated primary antibody was used, the cells were washed three times in cold FACS buffer and incubated in fluorescent secondary antibody diluted in FACS buffer on ice in the dark for 30 minutes. After the secondary antibody incubation or if a fluorescent-conjugated primary antibody was used, the cells were washed in cold FACS buffer three times and resuspended in cold FACS buffer. Right before each cell sample was run on the flow cytometer, Sytox Blue Dead Cell Stain (Invitrogen, S34857) was added to the sample to a final concentration of 1 μM. Then, 20,000-50,000 cells per sample were collected on a CytoFLEX Flow Cytometer (Beckman Coulter).

FlowJo software was used to analyze the FCS file of each sample. First, a dot plot was created for the side scatter height (SSC-H; y) and the forward scatter height (FSC-H; x) to create a “cell” gate, excluding small and large debris. From the cell gate, a dot plot was created for the side scatter height (SSC-H; y) and the side scatter area (SSC-A; x) to create a “single cell” gate, excluding the doublets. From the single cell gate, dot plot was created for the Pacific-Blue area (PB450-A; y) and the forward scatter area (FSC-A; x) to exclude dead cells, to create a “live cell” gate. From the live cell gate, the mean fluorescence intensity (e.g. APC-A or PE-A) was captured and used for downstream analysis.

To quantify antibody binding capacity (ABC), we utilized the Quantum^TM^ Simply Cellular® anti-Rat IgG (Bangs Laboratories, 817) kit following the manufacturer’s instructions. Briefly, microspheres were labeled with the same fluorescent secondary or fluorescent primary antibody used above at the same time as sample incubation. The stained microspheres were run by flow cytometry as described above to generate a standard curve capable of converting median fluorescence intensity to ABC values using the FREE QuickCal® v. 2.3 quantitative analysis program (Bangs Laboratories).

### Antibody Internalization Assay

Cells were plated in a 96-well plate (500 – 1000 cells per well) and allowed to seed for 48 hours, followed by the addition of the indicated drug and antibody. To monitor fluorescence, we used the Incucyte® Fabfluor-pH Antibody Labeling Reagent (Sartorius, 7722 (human), 4723 (Mouse)) following the manufacturer’s instructions. In brief, we mixed the primary antibody or ADC and Fabfluor-pH Antibody Labeling Reagent and diluted them in equal molar ratios in PBS and incubated the mixture at 37°C in the dark for 15 minutes. Following incubation, the indicated drug was added to the cells along with the antibody internalization reagent mixture. Plates were monitored using an incuyte live-cell imager to acquire brightfield and red fluorescent images of each well at 3-hour intervals for 72 hours. Internalization was quantified by normalizing the captured fluorescence intensity to the total cell area using Incucyte software and plotted on GraphPad PRISM.

### Crystal Violet Cell Growth Assay

Cells were seeded in 6-well plates at a low density (1000 – 5000 cells per well) 24 hours before the addition of the indicated drug. Media with and without drug was refreshed every 4 days thereafter. After 14 days of treatment, the media was aspirated, and cells were fixed in 10% buffered formalin (Fisher Scientific, SF-100-20) for 20 minutes. The formalin was aspirated and an equal volume of crystal violet solution (0.1% crystal violet in 10% ethanol) was added to each well for 20 minutes. After staining, the crystal violet solution was aspirated, and the plate was rinsed in running tap water until the excess crystal violet was removed. The plates were left to dry in the open air overnight and imaged on an Epson Perfection V600 Photo scanner. Images were analyzed in FIJI by setting a threshold to capture the crystal violet stain and measuring the percent area fraction of each well. The percent area was normalized to the vehicle control and plotted in GraphPad PRISM.

### Immunofluorescence microscopy

Cells were seeded onto ethanol-sterilized coverslips (Fisher, 1254583) in 12-well plates and treated with either DMSO or 100 nM MRTX1133 for 48 hours. Cells were washed twice with PBS and then fixed with 5% paraformaldehyde for 10 min at room temperature. Cells were washed twice with PBS, blocked with 1.5% goat serum for 1 hour, and washed in PBS three times. Cells were stained with anti-ALPP (Novus, NB110-3638, 1:250) for 1.5 hours on ice in a humidified chamber. Cells were washed three times with PBS and then stained with secondary antibody AlexaFluor488 (Invitrogen, A11001) for 1 hour at room temperature. Fluorescently labeled samples were then mounted with ProLong Gold Antifade Reagent with DAPI (CST, 8961S) on microscopy slides (Fisher, 12-550-143). All images were collected using a Zeiss AxioObserver with a Zeiss APOCHROMAT 63x oil objective and a photon multiplier tube detector. DAPI and ALPP were either excited at 405 nm (blue) or 488 nm (green) and emissions were collected. For each condition, 5 representative fields were captured for all samples.

### siRNA Transfection

Targeting constructs used were ON-TARGETplus 2.0 siRNA Dharmacon. Two distinct siRNAs targeting STAT3 were used: siSTAT-3-1 (UGAAAGUGGUAGAGAAUCU) and siSTAT3-2 (CGAUGGAGUACGUGCAGAA), and two distinct siRNAs targeting ALPP were used: siALPP-1 (CGGAAACGGUCCAGGCUAU) and siALPP-2 (AAAGGAAGUGUUAGACAAU). A control construct, Stealth RNAi™ siRNA Negative Control (siCTRL, Invitrogen, #12935300) was used for all knockdown experiments. Reverse transfections were performed for siRNA-mediated knockdown of indicated proteins by preparing a solution of siRNA and Lipfectamine RNAiMAX (Invitrogen, 13778150) and Opti-MEM Reduced Serum Medium (Gibco, 31985070). Equal volumes of siRNA diluted in Opti-MEM (final concentration of 2 nM for siSTAT3 and 10 nM for siALPP) and RNAiMAX diluted in Opti-MEM (final dilution of 1:1000) were mixed together and allowed to incubate for 30 min at room temperature. During this incubation, cells were trypsinized and seeded into plates. Following the incubation, the siRNA-RNAiMAX solution was added dropwise to the cell suspension and gently mixed.

### ALPP/ALPPL2 Serum Quantification

Quantitative measurement of ALPP/ALPPL2 in serum was performed using the Human Placental alkaline phosphatase SimpleStep ELISA® Kit (Abcam, #ab303758) as per the manufacturer’s protocol. Briefly, serum samples from mice were diluted 1:5 in Sample Diluent NS, and then 50 µL samples and standards were added to individual wells, followed by 50 µL of the Antibody Cocktail to each well. The plate was sealed and incubated for 1 hour at room temperature on a plate shaker set to 400 rpm. Following incubation, each well was washed three times with 350 µL 1X Wash Buffer PT. Next, 100 µL of TMB Development Solution was added to each well and allowed to incubate for 10 minutes in the dark on a plate shaker at 400 rpm. Next, 100 µL of Stop Solution was added to each well and placed on a plate shaker for 1 min to mix. Signal was quantified by recording the OD at 450 nm.

### Binding Affinity

High-binding 96-well MaxiSorp plates (Nunc) were coated overnight at 4 °C with human ALPPL2 (0.5 µg/mL in carbonate buffer: 8.4 g/L NaHCO_3_, 3.56 g/L Na_2_CO_3_). Plates were washed with phosphate-buffered solution (PBS) containing 0.05% Tween-20, blocked with 5% FCS in PBS/0.05% Tween-20 for 1 hour at 37 °C, and incubated overnight at 4 °C with serial dilutions of AT04 humanized antibodies (AT04-5A, AT04-13A, and AT04-23A) or isotype antibody. After washing, bound antibodies were detected with goat anti-human IgG–HRP (Jackson ImmunoResearch, 1:5000), developed with 3,3’,5,5’-tetramethylbenzidine (TMB) (Promega), stopped with 2N H_2_SO_4_, and absorbance at 450 nm was read on a Thermo Variolux plate reader. EC_50_ values were obtained by nonlinear curve fitting of absorbance versus log antibody concentration.

### Antigen Selectivity of AT04-26A

ELISAs were performed on high-binding 96-well MaxiSorp plates (Nunc). Antigens (recombinant human ALPI (CUSABIO), ALPL (Sino Biological), ALPP (Sino Biological), human ALPPL2 (Biointron), monkey ALPPL2 (Biointron), and mouse ALPG-His (Acro Biosystems) were diluted to 0.5 µg/mL in carbonate coating buffer (8.4 g/L NaHCO_3_, 3.56 g/L Na_2_CO_3_, pH 9.5) and 50 µL/well were incubated overnight at 4 °C. Plates were washed with PBS (3x) containing 0.05% Tween-20 (PBS-T), rinsed with PBS, and blocked with 5% fetal calf serum (FCS) in PBS-T (100 µL/well, 1 hour, 37 °C). Human IgG1 antibody AT04-23A, and isotype antibody were prepared at micromolar stock concentrations and serially diluted 5-fold in assay buffer to yield a final range of 20– 0.00128 nM. Fifty microliters of each dilution were added per well and incubated for 1.5– 3 hours at 37°C, followed by washing. HRP-conjugated goat anti-human IgG (1:5000 in assay buffer) was added (50 µL/well, 1 hour, 37 °C), and plates were washed before adding TMB substrate. Reactions were stopped with an equal volume of stop solution, and absorbance at 450 nm was measured using a Varioskan LUX reader. EC_50_ values were determined by nonlinear regression using a four-parameter logistic model.

### Surface Plasmon Resonance (SPR)

A Biocore T200 (Cytiva) was used to determine the affinity of AT04-26A to human ALPPL2. The amine coupling approach was used to immobilize anti-human IgG Fc proteins onto a CM5 chip at 10 mL/min in 10 mM sodium acetate buffer (pH 5.0). The sensor surface was activated with a 7 min injection of 50 mM NHS and 200 mM EDC, followed by the injection of 45 μg/mL Anti-Human IgG Fc proteins to achieve a target level of 19687.4 RU. The surface was then blocked with 1 M ethanolamine (pH 8.5). A solution of antibody AT04-26A (1 mg/mL) in HBS-N (10 mM HEPES, 150 mM NaCl, pH 7.4) running buffer injected at 10 mL/min for 60 seconds to allow capture on the surface of the CM5-conjugates anti-human IgG Fc chip. Recombinant ALPPL2-FLAG antigen was tested at five concentrations (12.3, 37, 111, 333, and 1000 nM) with a flow rate of 10 mL/min and a binding time of 60 seconds to determine the association and dissociation rate constants. The experiment was performed in triplicate with response signals plotted against time. The results were analysed with the BIAcore T200 software visualized with GraphPad Prism.

### Bioconjugation of AT04-26A

AT2604 was generated by site-specific conjugation of antibody AT04-26A to a vc-PABC– MMAE linker–payload. Briefly, antibody was buffer-exchanged into 50 mM Tris-HCl, 2 mM EDTA, pH 8.0, and reduced with 200 molar equivalents of dithiothreitol (DTT) at 22 °C overnight to fully reduce engineered cysteines. The reduced antibody was then buffer-exchanged into 50 mM Tris-HCl, 150 mM NaCl, 2 mM EDTA, pH 7.5, followed by addition of 20 molar equivalents dehydroascorbic acid (DHAA) and incubation for 0.5 h at 22 °C to re-oxidize native disulfide bonds while preserving engineered thiols for conjugation.

For conjugation, 6 molar equivalents of mc-vc-PABC-MMAE (MedChemExpress) were added to the thiol-containing antibody and the reaction mixture was incubated for 2 hours at 22 °C with gentle agitation to allow thiol–maleimide coupling. Unconjugated linker– payload was removed by desalting using a Zeba^TM^ spin column, and the resulting ADC was buffer-exchanged into 50 mM histidine–acetate, pH 5.5, by repeated concentration– dilution (4–6 cycles) using a 30 kDa MWCO polyethersulfone ultrafiltration device, then stored at −80 °C.

### Determination of Drug to Antibody Ratio (DAR)

Drug–to–antibody ratio (DAR) of cysteine-conjugated mc-vc-PABC-MMAE antibody–drug conjugates were determined by hydrophobic interaction chromatography (HIC) followed by intact mass spectrometry. ADC samples were analyzed on a butyl HIC column (MAbPac HIC-Butyl, 4.6 × 100 mm, ThermoFisher) operated at 25–30 °C with UV detection at 280 nm. Mobile phase A consisted of 1.5M ammonium sulfate, 50 mM dipotassium phosphate, pH 7.0; mobile phase B consisted of 50 mM dipotassium phosphate, pH 7.0, with 25% isopropanol (v/v). Samples were injected and eluted at 1 mL/min using a linear gradient from 100% A to 100% B over 20–30 min, followed by a high-B wash and re-equilibration at initial conditions. Under these conditions, vc-MMAE ADCs eluted as discrete peaks corresponding to species with 0, 2, 4, 6, and 8 drugs per antibody.

The average DAR was calculated from the relative UV peak areas for each conjugation species using DAR = Σ(n·A)/ΣA, where n is the drug load and A is its integrated peak area. To confirm species assignment and average DAR, selected samples were analyzed by intact LC–MS under native-compatible conditions (e.g., desalting on a short reversed-phase column with volatile ammonium acetate buffer), and deconvoluted mass spectra were used to verify the observed mass of each DAR species.

### In vitro cytotoxicity

NCI-N87 (ATCC) gastric carcinoma cells were cultured in RPMI-1640 (ThermoFisher) supplemented with heat inactivated FBS 10%, at 37 °C under 5% CO_2_. Cells at logarithmic growth were harvested and seeded at 3000 cells per well in 96-well plates in 90 μL culture media then incubated overnight. AT2604 and isotype ADCstock solutions were diluted with media to generate 4-fold serial dilutions The next day, 10 μL of each dilution was added per well and cells were incubated for five days (120 hours). At endpoint, plates and CellTiter-Glo 2.0 (Promega) were equilibrated to room temperature for 30 min, 50 μL reagent was added directly to each well, plates were shaken for 5 min, and luminescence was read on a Varioskan Lux (ThermoFisher) microplate reader. Curves and CC_50_ values were generated in GraphPad Prism Version 9.4.1, using non-linear regression, curve fit for inhibitor vs response.

### Antibody-Dependent Cellular Cytotoxicity (ADCC) activity of AT04-26A

PBMC-mediated cytotoxicity assay using impedance-based real-time cell analysis (RTCA) was performed over two days. Peripheral blood mononuclear cells (PBMCs) were used as effector cells, and adherent target cells were seeded onto electronic microtiter plates for continuous monitoring of cell index.

Frozen PBMCs were thawed and transferred into RPMI 1640 (Gibco) supplemented with 10% fetal bovine serum (FBS), then incubated overnight at 37 °C under 5% CO₂. Target cells were harvested, counted, and resuspended in assay buffer (RPMI 1640 containing 10% FBS) at a concentration of 200,000 cells/mL. For plate preparation, 50 µL of 10% FBS–RPMI 1640 medium was first added to each well of the electronic 96-well plate (E-plate) to equilibrate the sensors, after which 50 µL of the target-cell suspension (corresponding to 10000 cells per well) was seeded. The plate was allowed to equilibrate for 30 min at room temperature, and then incubated overnight at 37 °C under 5% CO₂.

The following day PBMCs were harvested, the supernatant was discarded, and cells were resuspended in culture medium and adjusted to 4×10^6^ cells/mL. Next, 50 µL was added to each well containing target cells to achieve an effector-to-target (E:T) ratio of 20:1. Test antibodies in culture medium were added at 50 µL per well to reach the desired final concentrations. Plates were immediately placed in an xCELLigence Real Time Cell Analyzer (Agilent) and cell index was monitored every 15 min for 6–24 hours. Cytotoxicity was quantified based on changes in cell index, and percentage cytotoxicity was calculated using the following equation: cytotoxicity (%) = (1 - sample/isotype control) x 100

### Immunohistochemical (IHC) staining

For IHC staining performed in Figures 5F-G, formalin-fixed paraffin-embedded (FFPE) tumor tissue microarrays (TMAs) were developed by Crown Bioscience. Both GA2419 and GA6203 are derived from gastric cancer patients with FGFR2 and MYC amplified, respectively. IHC staining was conducted following a standard protocol. Briefly, paraffin sections were deparaffinized with xylene and rehydrated through a graded ethanol series, followed by antigen retrieval using EDTA buffer (pH 6). After cooling, endogenous peroxidase was blocked with 3% hydrogen peroxide. Sections were rinsed with tris-buffered saline with Tween 20 (TBST) and blocked with 5% BSA to reduce non-specific binding. The sections were incubated with primary antibody against ALPPL2 (rabbit polyclonal antibody from Origen, TA308201, diluted 1:400) at room temperature for 1 hour, then washed and incubated with HRP-conjugated goat anti-rabbit secondary antibody. After further TBST washing, 3,3’-diaminobenzidine (DAB) was applied for color development, and the reaction was terminated with purified water. Nuclei were counterstained with hematoxylin. Finally, sections were dehydrated with absolute ethanol, cleared in xylene and mounted with neutral balsam. Immunohistochemical staining results were evaluated semi-quantitatively. All slides were scanned for subsequent qualitative and semi-quantitative analysis. The IHC staining results of ALPPL2 were evaluated using a standardized semi-quantitative scoring system based on positive cell percentage and staining intensity. According to the overall staining degree and positive cell proportion, the staining results were classified into four grades: negative (-, no positive staining), weakly positive (+, pale yellow staining in partial cells), moderately positive (++, uniform brown staining in most cells), and strongly positive (+++, intense dark brown staining in almost all cells). The grading results were used to evaluate the low and high expression levels of ALPPL2 for subsequent statistical analysis. H-score was determined using the following formula: H-score = (1 x %cells 1+) + (2 x %cells 2+) + (3 x %cells 3+).

For IHC staining in Figure 6G and Supplementary Figure S8E, slides were placed in oven at 60 °C for 20 min and allowed to cool for 5 min at room temperature. Slides were submerged in xylene for 5 minutes two times, rehydrated by submerging in decreasing alcohol concentrations for 5 minutes each (100%, 100%, 95%, and 70%), and submerged in DI water for 3 minutes. Antigen retrieval was performed by submerging slides in 1x sodium citrate pH 6 solution (Agilent Dako, S2369) and heated in a pressure cooker under low pressure for 30 minutes. Slides were removed and allowed to cool to room temperature and then washed in 1x TBS twice for 3 minutes each. Upon using a PAP pen to draw a hydrophobic barrier around tissue sections, samples were incubated with 3% hydrogen peroxide solution in TBS for 10 min at room temperature. Samples were washed in TBS for 1 min three times. Tissues were blocked with goat serum in TBST for 1 hour at room temperature, and washed again. Slides were incubated with indicated primary antibody (ALPP/ALPPL2, Abcam Ab133602, 1:250; p-ERK (T202/Y204) Cell Signaling Technology, 4370, 1:400) in a humidified chamber at 4 °C overnight. Samples were washed in TBST, and SignalStain® Boost IHC Detection Reagent (Cell Signaling Technology, #8114) was added for 30 minutes at room temperature. Samples were washed with TBST and then incubated with DAB for 2-5 minutes, then quenched in Milli-Q water. Counterstaining with hematoxylin was performed for 2 min, and slides were washed under running water until water ran clear. Samples were destained in 70% EtOH 0.25% HCl (250 mL EtOH 70% + 625 uL HCl 12N) for 7 quick dunks, and rinsed in Milli-Q water for 3 minutes. Dehydration of tissue was performed using increasing alcohol concentrations for 3 min each (70%, 95%, 100%, and 100%), then cleared in xylene twice for 5 min each. Slides were allowed to dry overnight and then mounted and cover slipped. Slides were allowed to dry overnight and imaged on an Akoya PhenoImager HT.

To quantify IHC staining, whole slide scans of ALPP and pERK stained tumor tissues were analyzed in Fiji. For each tissue section, five representative viable tumor fields (excluding necrotic tissue) were selected. Each field was color deconvoluted to isolate the DAB channel, and mean pixel intensity was measured and converted to optical density using OD = log_10_(background / mean intensity). The five fields were averaged and comparisons were made between pooled vehicle and RMC-6236 treated tumors (circles = 3 days, triangles = 7 days).

## Acknowledgements

S.Liu, B.K.S. Yeung, and B. Zou would like to acknowledge past Axcynsis Therapeutics team members who made contributions to the discovery and development of AT2604. We thank the Dana-Farber/Harvard Cancer Center in Boston, MA, for the use of the Specialized Histopathology Core, which provided histology and immunohistochemistry service. Dana-Farber/Harvard Cancer Center is supported in part by an NCI Cancer Center Support Grant # NIH 5 P30 CA06516. J.A. Paulo is funded by R35 GM156406.

C.A.S was supported by NCI P50CA196510. A.J. Aguirre and J.D. Mancias were funded by the Hale Family Center for Pancreatic Cancer Research at Dana-Farber Cancer Institute. This Break *Through* Cancer KRAS TeamLab study reflects a cross-institutional collaboration, supported (in part) by the Break *Through* Cancer and the Lustgarten Foundation for Pancreatic Cancer Research (J.D. Mancias, A.J. Aguirre, T.E. Oni and T.P. Heffernan). A.J. Aguirre is also funded by the Pancreatic Cancer Action Network and NIH R01 CA276268. J.D. Mancias is also funded by NIH R01 CA301542.

## Authors’ Contributions

J.M. DeLiberty Conceptualization, formal analysis, data curation, investigation, visualization, writing–original draft, writing–review and editing, supervision, methodology, project administration.

A. F. Kong Conceptualization, formal analysis, data curation, investigation, visualization, writing–original draft, writing–review and editing, supervision, methodology, project administration.

B. Q. Yu Conceptualization, formal analysis, data curation, investigation, visualization, writing–review and editing, supervision, methodology, project administration.

C. N. Sindoni Investigation, formal analysis, methodology, writing–review and editing, project administration.

D. J. Lin Investigation, formal analysis, validation, writing–review and editing.

E. K. Kang Investigation, validation, writing–review and editing.

E.Y. Kim Investigation, validation, writing–review and editing.

S. Chen Investigation, validation, writing–review and editing.

A. Padhye Investigation, formal analysis, data curation, supervision, writing–review and editing.

O. Coker Formal analysis, data curation, investigation, visualization, writing–review and editing.

J.H. Choe Formal analysis, data curation, investigation, visualization, writing–review and editing.

J.A. Paulo Investigation, methodology, writing–review and editing.

M. Shin Investigation, methodology, writing–review and editing.

L. Hendrianto Investigation, methodology, writing–review and editing.

K. Takahashi Investigation, formal analysis, methodology, supervision, writing–review and editing.

J. Varga-Wiles Investigation, writing–review and editing.

B.D. Parent Investigation, writing–review and editing.

Z. Li Investigation, writing–review and editing.

S. Shubitidze Investigation, writing–review and editing.

T.E. Oni Conceptualization, investigation, writing–review and editing, supervision, project administration, funding acquisition.

D.H. Peng Formal analysis, investigation, writing–review and editing, supervision, project administration.

T.P. Heffernan Investigation, writing–review and editing, supervision, project administration.

S. Liu Conceptualization, formal analysis, data curation, investigation, writing–review and editing, supervision, project administration.

B.K.S. Yeung Conceptualization, formal analysis, data curation, investigation, writing– original draft, writing–review and editing, supervision, methodology, project administration.

B. Zou Conceptualization, formal analysis, data curation, investigation, writing–review and editing, supervision, methodology, project administration, funding acquisition.

C.A. Stalnecker Conceptualization, formal analysis, data curation, investigation, visualization, writing–original draft, writing–review and editing, supervision, methodology, project administration, funding acquisition.

A.J. Aguirre Conceptualization, formal analysis, data curation, investigation, visualization, writing–original draft, writing–review and editing, supervision, methodology, project administration, funding acquisition.

J.D. Mancias Conceptualization, formal analysis, data curation, investigation, visualization, writing–original draft, writing–review and editing, supervision, methodology, project administration, funding acquisition.

## SUPPLEMENTARY FIGURE LEGENDS

**Supplementary Fig. S1. PDAC model response to MRTX1133 treatment. A.** Relative viability of PDAC cell lines treated with a dose range of MRTX1133 for 5 days. **B.** Protein levels of phospho-ERK (T202/Y204) and total ERK following treatment of KRASi for 6 h and 2 d (MRTX1133, 100 nM) or for 7 d (MRTX1133, 20 nM) as measured by immunoblot. Par = Parental, RST = MRTX1133-Resistant (cultured in 1 µM MRTX1133). **C**. Relative viability of parental (-Par) and MRTX1133-resistant (-MR) isogenic PDAC cell lines treated with a dose range of MRTX1133 for 5 days. **D.** Relative tumor volume fold change of CDXs treated with either vehicle or 30 mg/kg MRTX1133 twice daily for 28 days. Measurements represent endpoint measurements relative to day 0. Statistical significance was determined by Welch’s t test.

**Supplementary Fig. S2. Baseline characterization and global proteomic variance across temporal KRAS inhibition models. A.** Principal component analysis (PCA) of global proteomic differential expression across *in vitro* cell lines. **B.** Pearson correlation matrix comparing the differential expression profiles of the *in vitro* proteomic datasets. **C.** Quantification of significantly downregulated (adj. *p* < 0.05, log_2_ fold-change < −0.5, left) and upregulated (log_2_ fold-change > 0.5, right) essential proteins (DepMap Chronos score < −0.75) across the AsPC-1, Panc02.03, PK-45H, and SUIT-2 cell lines at the indicated timepoints. **D.** GSEA of canonical KEGG and REACTOME gene sets evaluated in *in vitro* temporal models and *in vivo* xenografts (naïve vs. acquired resistance).

**Supplementary Fig. S3. Pathway clustering and individual subtype marker dynamics in response to KRAS inhibition. A.** Szymkiewicz-Simpson overlap matrix of the top 25 significantly differentially regulated KEGG and REACTOME pathways at each timepoint, colored by temporal cluster. The optimal clustering method and number of clusters (k=4) were determined by agglomerative coefficient and gap statistic analyses. The top heatmap displays the corresponding GSEA net enrichment score (NES) for each pathway across the temporal timepoints. **B.** Log_2_(fold-change) values of individual basal or classical subtype marker proteins. Data represent aggregate global proteomics across *in vitro* acute KRASi (2 days), *in vivo* models, and matched acute/sustained *in vitro* models (AsPC-1, Panc02.03, PK-45H, and SUIT-2). Statistical significance was evaluated using a one-sample t-test comparing the mean log_2_(fold-change) to 0.

**Supplementary Fig. S4. Phosphoproteome and kinome rewiring, regulatory site dynamics, and individual temporal kinase-substrate networks following KRAS inhibition. A, B.** Principal component analysis (PCA) (**A**) and Pearson correlation matrix (**B**) of differentially regulated phosphosites across *in vitro* temporal models. **C.** Quantification of significantly down- (log_2_ fold-change < −0.5, left) and up-regulated (log_2_ fold-change > 0.5, right) regulatory phosphosites across timepoints. **D.** Pearson correlation analysis of inferred kinase activities (motif frequency factors) across the *in vitro* time course. **E.** Inferred kinase activities across timepoints represented by log_2_(frequency factor) versus significance (-log_10_ adj. *p*-value), with top dysregulated kinases labeled. **F–H.** Individual kinase-substrate networks capturing adaptations at 2 days (**F**), 7 days (**G**), and acquired resistance (**H**). Network topologies, node characteristics (DepMap Chronos scores), shapes (diamonds for kinases, circles for substrates), and edge weightings mirror the specifications described in Fig. 2C.

**Supplementary Fig. S5. Integration of global and BAMX-enriched proteomics expands the KRAS-regulated surfaceome and characterizes candidate ADC targets. A.** Aggregate volcano plots of BAMX-enriched surface proteins in four longitudinally tracked PDAC cell lines at 7 days of KRASi and acquired resistance. Clinically investigated ADC targets are plotted below (red: significantly upregulated, adj. *p* < 0.05, log2FC > 0.5; blue: significantly downregulated, log2FC < −0.5). **B.** Venn diagrams depicting the overlap of surface-annotated proteins detected via global versus BAMX-enriched proteomics at 2 days, 7 days, and acquired resistance. **C.** Correlation analysis of surface protein differential expression (log_2_ fold-change) captured by global proteomics versus BAMX enrichment at 7 days and resistance. Green dots indicate shared proteins; red (top) and blue (right) dots represent proteins unique to global or BAMX datasets, respectively. Significance thresholds (adj. *p* < 0.05, |log2FC| > 0.5) are denoted for uniquely detected proteins. **D.** Histograms quantifying the number of significantly upregulated clinical ADC targets (adj. *p* < 0.05, log_2_ fold-change > 0) captured by each individual modality compared to the integrated dataset combined by Stouffer’s weighted z-score. **E.** Line plot showing the coefficient weights (x-axis) assigned to each multi-omic feature (y-axis) utilized to train the machine learning model. Features driving a higher “ADC score” exhibit positive coefficients; these include surface annotation, experimental KRASi-induced upregulation (surfaceome proteomics log_2_FC, global proteomics log_2_FC, and RNAseq log_2_FC), tumor-enriched expression metrics (PDAC IHC score, TCGA PAAD RNA expression, and CPTAC PAAD tumor-normal log_2_FC), and prognostic features (HPA PDAC prognostic factor). Conversely, features representing normal tissue expression (normal tissue IHC score and normal tissue RNA expression) exhibit negative coefficients. **F.** Multi-omic feature heatmap of prioritized ADC candidates, including the top 10 highest-scoring proteins, bottom 5 lowest-scoring proteins, and selected resistance-associated receptors (e.g., EGFR, AXL, MET). Columns denote the specific training features utilized for each protein, including temporal KRAS-regulated dynamics (integrated proteomics at 2 and 7 days; RNA-seq at 24 hours), expression disparities in normal versus PDAC tissues (CPTAC, TCGA, HPA IHC), prognostic significance, clinical ADC status, and the final computed KRASi ADC Target Score.

**Supplementary Fig. S6. Validation of non-ALPP/ALPPL2 surface targets, published ALPP expression, and siRNA-mediated knockdown of ALPP. A.** Mean fluorescence intensity of indicated surface targets following DMSO treatment, 2 day KRASi (MRTX1133, 100 nM), or 7 day KRASi (MRTX1133, 20 nM). Each data point represents an independent experiment, error bars denote mean ± s.d., n = 3 independent repeats. Statistical significance was determined by two-way ANOVA. **B.** Expression of ALPP in indicated healthy tissue from GTEx dataset. **C.** Internalization of the indicated antibody (10 nM), quantified by total integrated density by Incucyte live-cell imaging. Panc 02.03 cells were treated with and without KRASi (MRTX1133; 100 nM). **D.** *ALPP* and *ALPG* transcript levels increase in response to MRTX1133 across timepoints in cell culture and in vivo models of PDAC (data from Hallin et al. Nature Medicine 2022; GSE201412)(40). *In vivo* AsPC-1 cells were treated with 100 nM MRTX1133 or DMSO for indicated timepoints. *In vivo* AsPC-1 xenograft mice received 30 mg/kg MRTX1133 BID for either 1.5 days (three doses every 12 hours and then collected 6 or 24 hours after the final dose, AsPC1 6h *in vivo* or AsPC1 1.5d *in vivo*, respectively) or 7 days. **E.** ALPP/ALPPL2 protein levels in AsPC-1 and PANC-1 treated with the indicated siRNA for 6 days were determined by western blotting. **F.** Relative cell viability of AsPC-1 and PANC-1 cells following siRNA-mediated ALPP knockdown and treatment with either vehicle or a dose-response of MRTX1133 for 5 days. *P<0.05, **P<0.01, ***P<0.001, ****P<0.0001; ns, not significant.

**Supplementary Fig. S7. Characterization of AT2604. A.** Antibody-dependent cellular cytotoxicity (ADCC) of AT04-26A is abrogated compared to the Fc-competent, AT04-27A antibody. Statistical significance was determined by two-way ANOVA. **B.** Hydrophobic Interaction Chromatography (HIC) trace showing high DAR 4 homogeneity after the conjugation step with vc-MMAE. **C.** Average body weight of mice enrolled in the study described in **Fig. 5E**. Arrows denote ADC treatment days, **D.** Average body weight of mice enrolled in the study described in **Fig. 5F**. Arrows denote ADC treatment days, **E.** Average body weight of mice enrolled in the study described in **Fig. 5G**. Arrows denote ADC treatment days. **F**. Representative histograms of ALPP/ALPPL2 expression in PDAC cells treated with DMSO, KRASi for 2 days (MRTX1133; 100 nM for AsPC-1, SW1990, Panc 02.03, HPAC, and PK-45H; 1 µM for PANC-1), or KRASi for 7 days (MRTX1133; 20 nM for AsPC-1, SW1990, Panc 02.03, HPAC, and PK-45H; 1 µM for PANC-1). Histograms are representative of three independent experiments. **G**. Quantified ALPP/ALPPL2 surface expression on OVCAR-5 and SNU-8 ovarian cancer cell lines treated with either 10 nM RMC-6236 or 100 nM MRTX1133, respectively, for 2 days as measured by flow cytometry. **H**. Relative *ALPP/ALPPL2* mRNA in SNU-8 and H1373 cell lines following treatment with KRASi (SNU-8: MRTX1133, 100 nM; H1373: adagrasib, 100 nM) or JAKi (Cucurbitacin I, 250 nM) alone or in combination. Statistical significance was determined by one-way ANOVA. *P<0.05, **P<0.01, ***P<0.001, ****P<0.0001; ns, not significant.

**Supplementary Fig. S8. AT2604 antibody characterization and combination of RASi and ALPP/ALPPL2 ADC leads to sustained tumor growth inhibition in PDAC models. A**. Internalization of the anti-AT2604 or AT-00-11A-MMAE or their parent antibodies (10 nM), quantified by total integrated density by Incucyte live-cell imaging. PANC-1 and HPAC cells were treated with and without KRASi (MRTX1133; 1 µM for PANC-1 and 10 nM for HPAC). Statistical significance was determined by one-way ANOVA. **B.** Relative cell viability of indicated PDAC cell lines treated with MMAE for 5 days. **C.** Relative *ALPP/ALPPL2* mRNA from two PDAC PDOs treated with 100 nM RMC-6236 for 2 days, corresponding to samples in **Fig. 6E**. **D.** Top, relative viability of PDAC PDOs treated with a dose range of RMC-6236 alone or in combination with increasing doses of AT-00-11A-MMAE for 6 days. Each data point represents the mean of triplicates ± s.d.; viability curves shown are representative of three independent experiments. Bottom, Highest Single Agent synergy score values and heatmaps from data represented above. Tiles in heat map represent HSA synergy scores (red = synergy, white = additivity, green = antagonism). **E**. Mean optical density (OD) quantified by IHC staining from samples in **Fig. 6G**. For each tissue section, five representative viable tumor fields were selected, and mean pixel intensity was converted to OD and averaged (circles = 3 days, triangles = 7 days). Statistics were determined by Welch’s t test. **F.** Individual tumor growth curves for mice enrolled in study as described in **Fig. 6I**. Arrows denote ADC treatment days, and the gray box denotes the dose schedule of 10 mg/kg RMC-6236 PO QDx14 (or vehicle). **G.** Average body weight of mice enrolled in study as described in **Fig. 6I** and **Supp. Fig. S6F**. Arrows denote ADC treatment days, and the gray box denotes the dose schedule of 10 mg/kg RMC-6236 PO QDx14 (or vehicle). *P<0.05, **P<0.01, ***P<0.001, ****P<0.0001; ns, not significant.

## Supplemental Tables

**Supplemental Table S1**: Metadata for proteomics searches

**Supplemental Table S2**: Differential expression of global proteomics

**Supplemental Table S3**: Gene set enrichment results for global proteomics data

**Supplemental Table S4**: Differential expression results for phosphoproteomics data

**Supplemental Table S5**: Kinase enrichment analyses

**Supplemental Table S6**: Differential expression results for surface-enriched proteomics

**Supplemental Table S7**: Integrated proteomics and surface-enriched proteomics datasets

**Supplemental Table S8**: Features used to train ridge logistic model

