## Supplementary Figures for "KRAS inhibition drives adaptive surfaceome remodeling and reveals ALPP/ALPPL2 as a targetable vulnerability in pancreatic cancer"

### Supplementary Figure S1

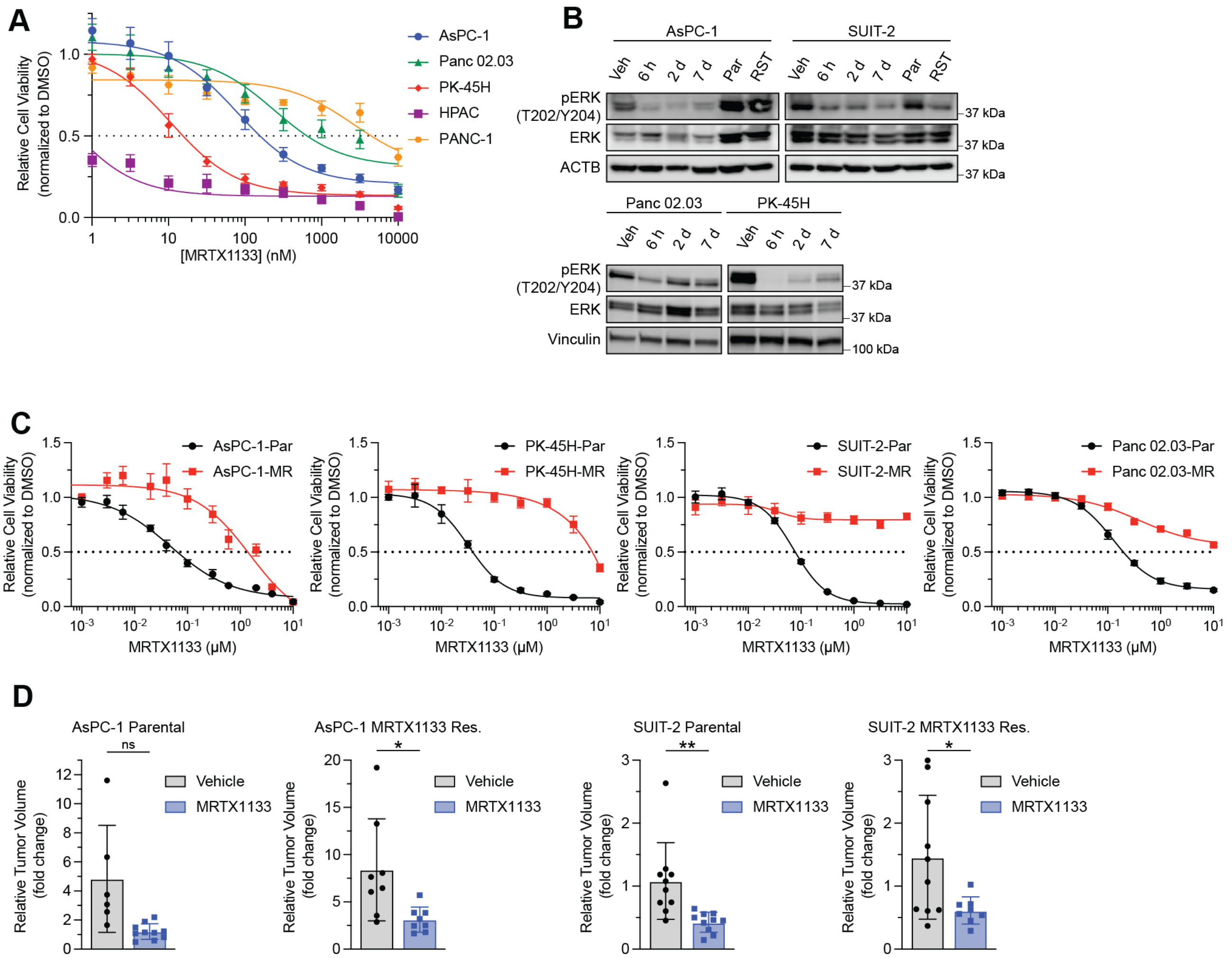

### Supplementary Figure S2

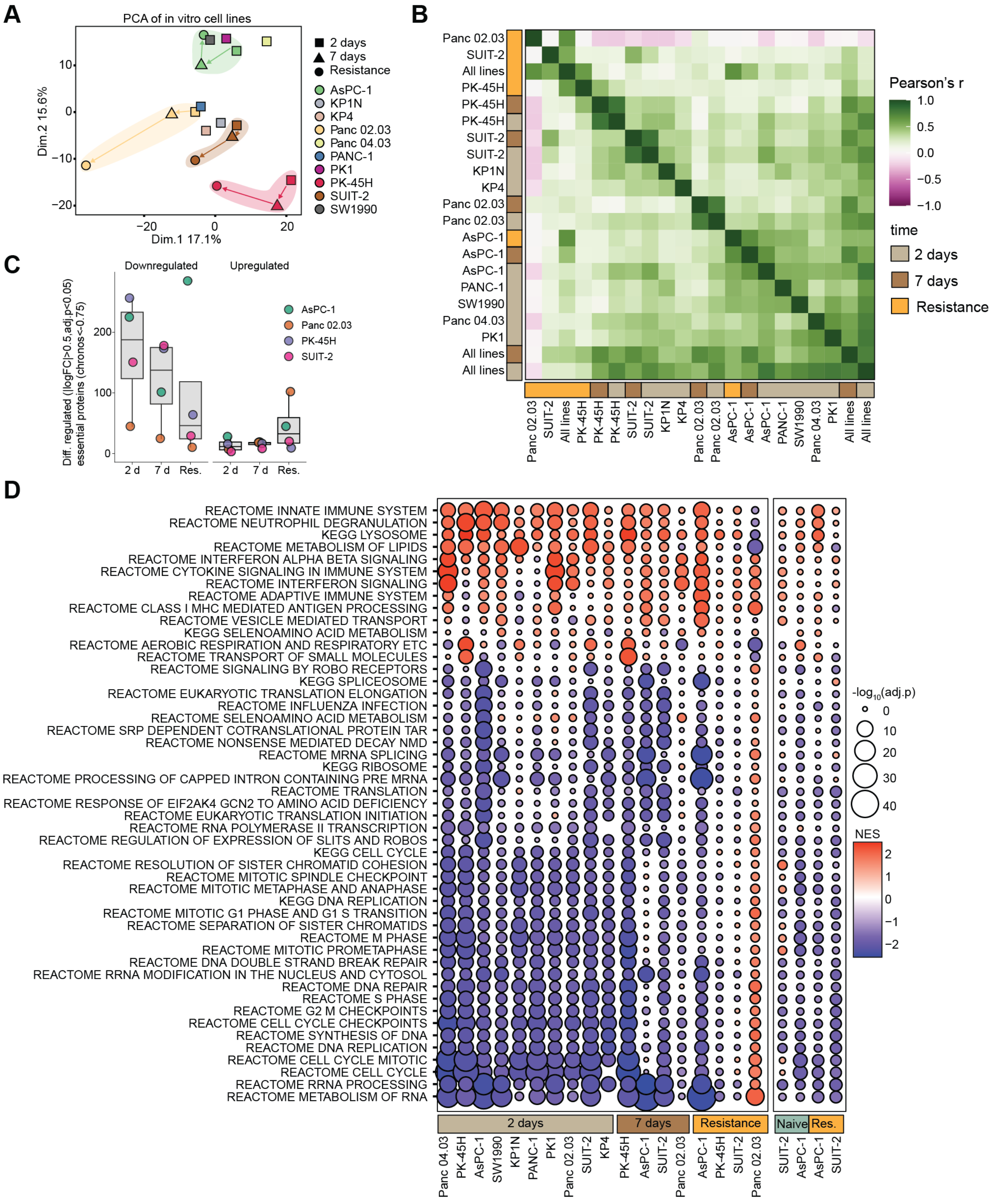

### Supplementary Figure S3

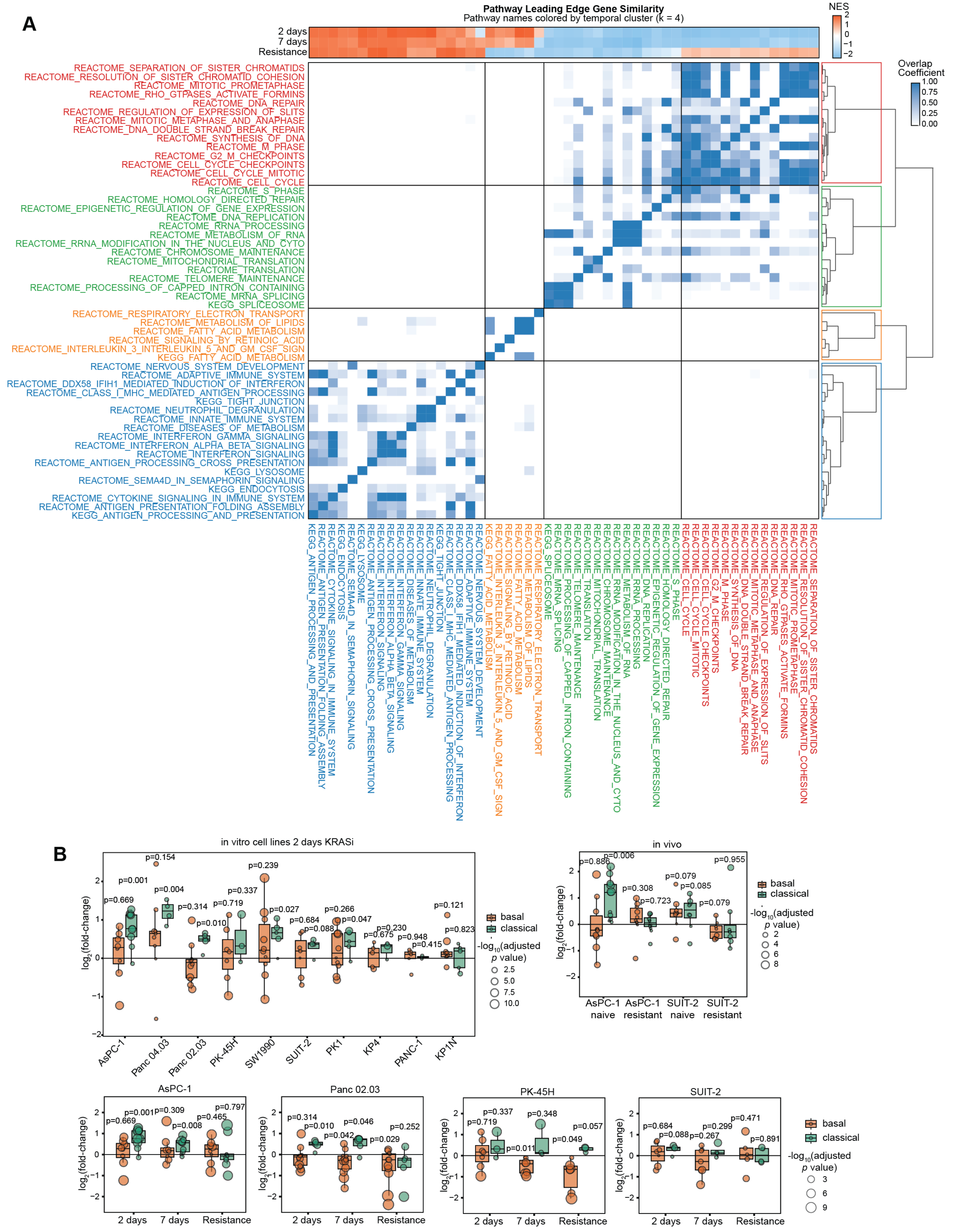

#### Supplementary Figure S4

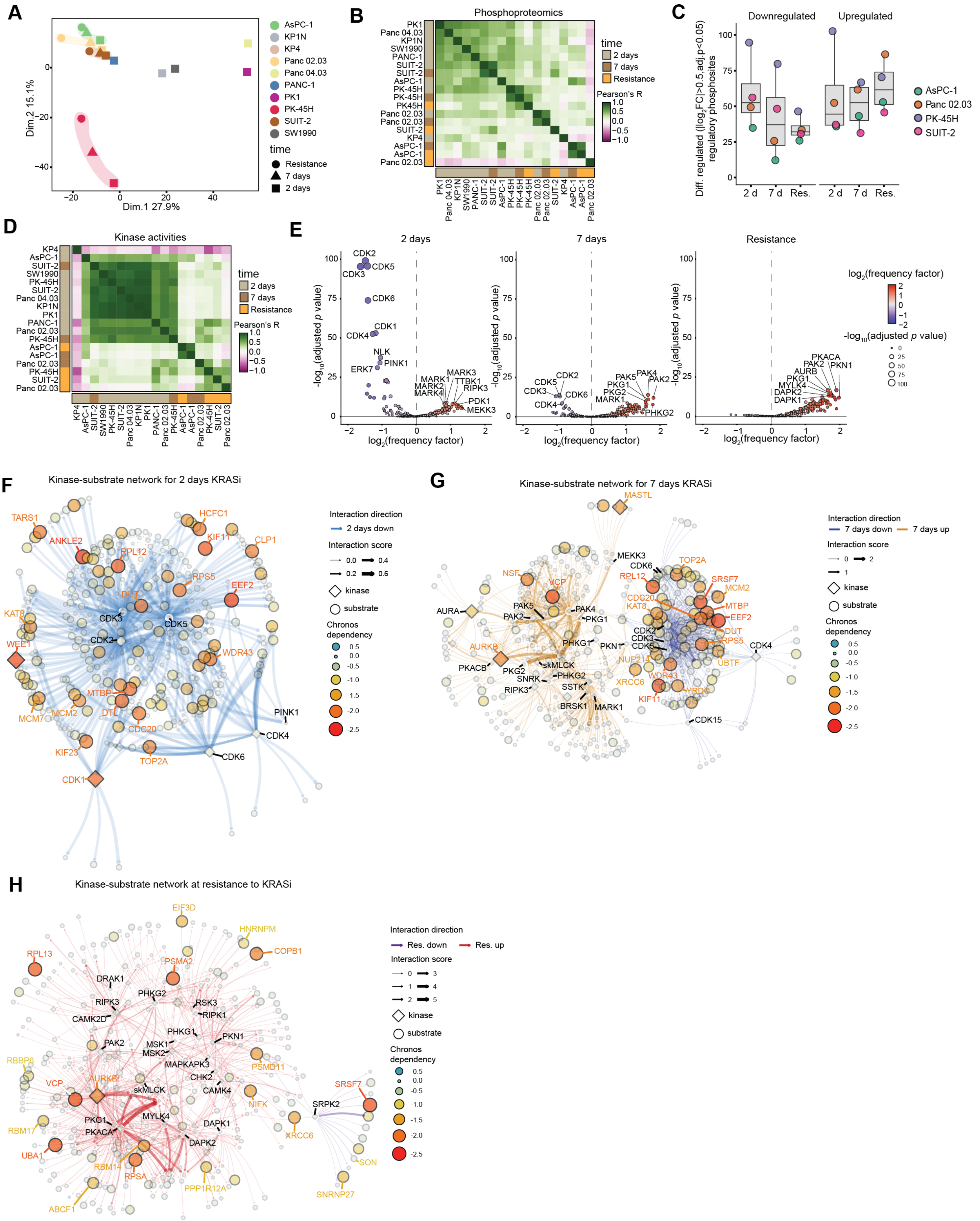

### Supplementary Figure S5

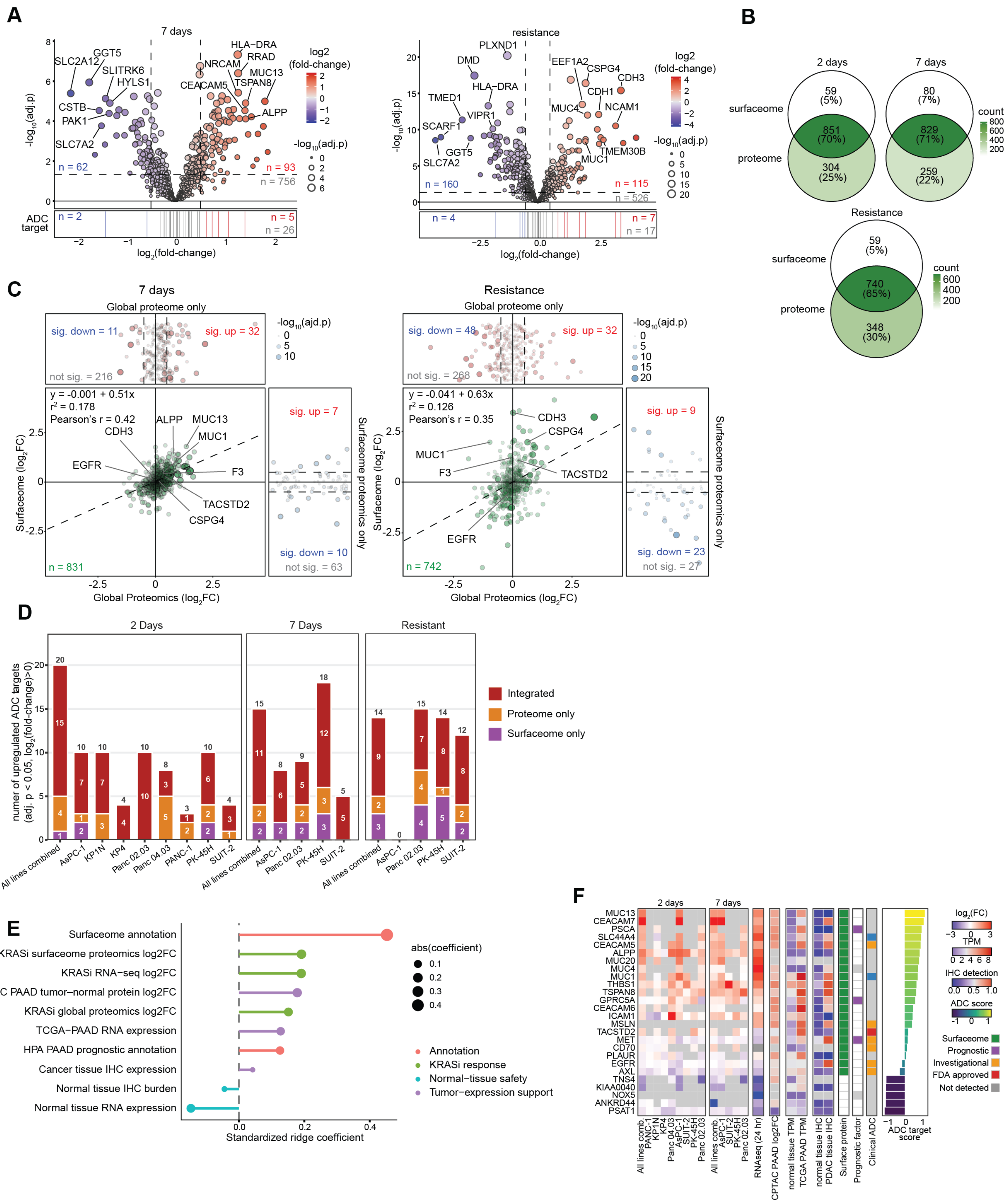

### Supplementary Figure S6

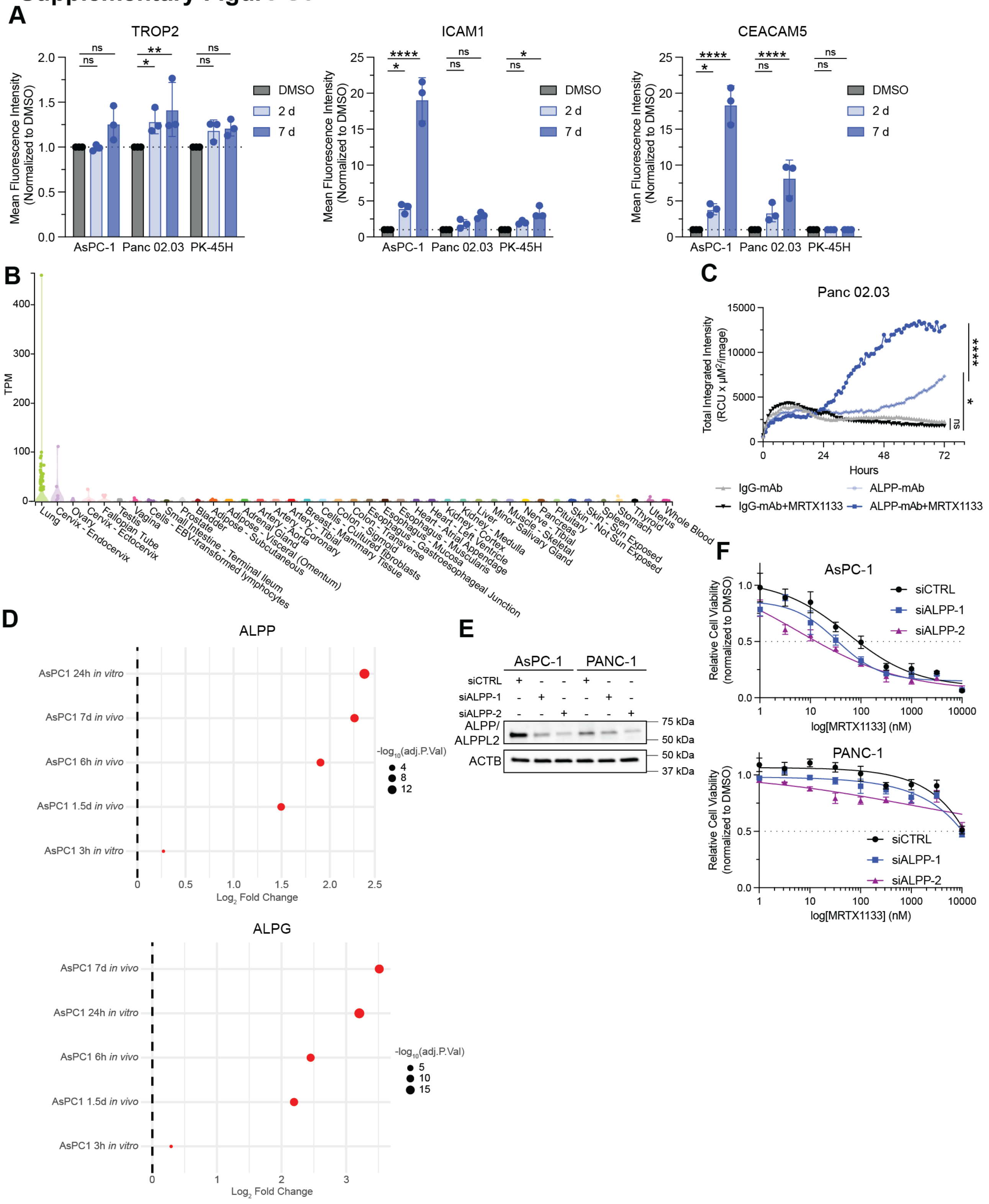

#### Supplementary Figure S7

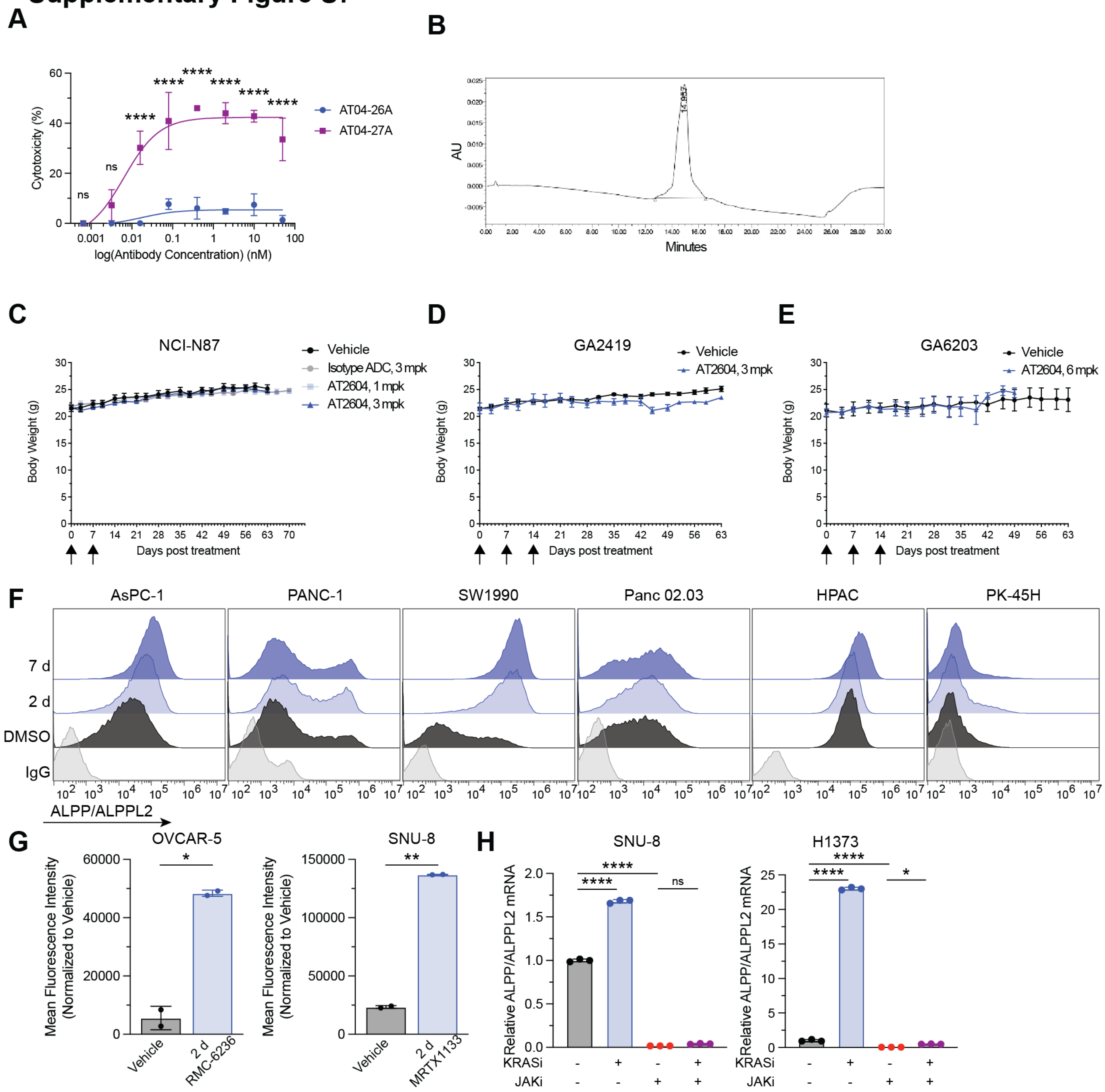

### Supplementary Figure S8

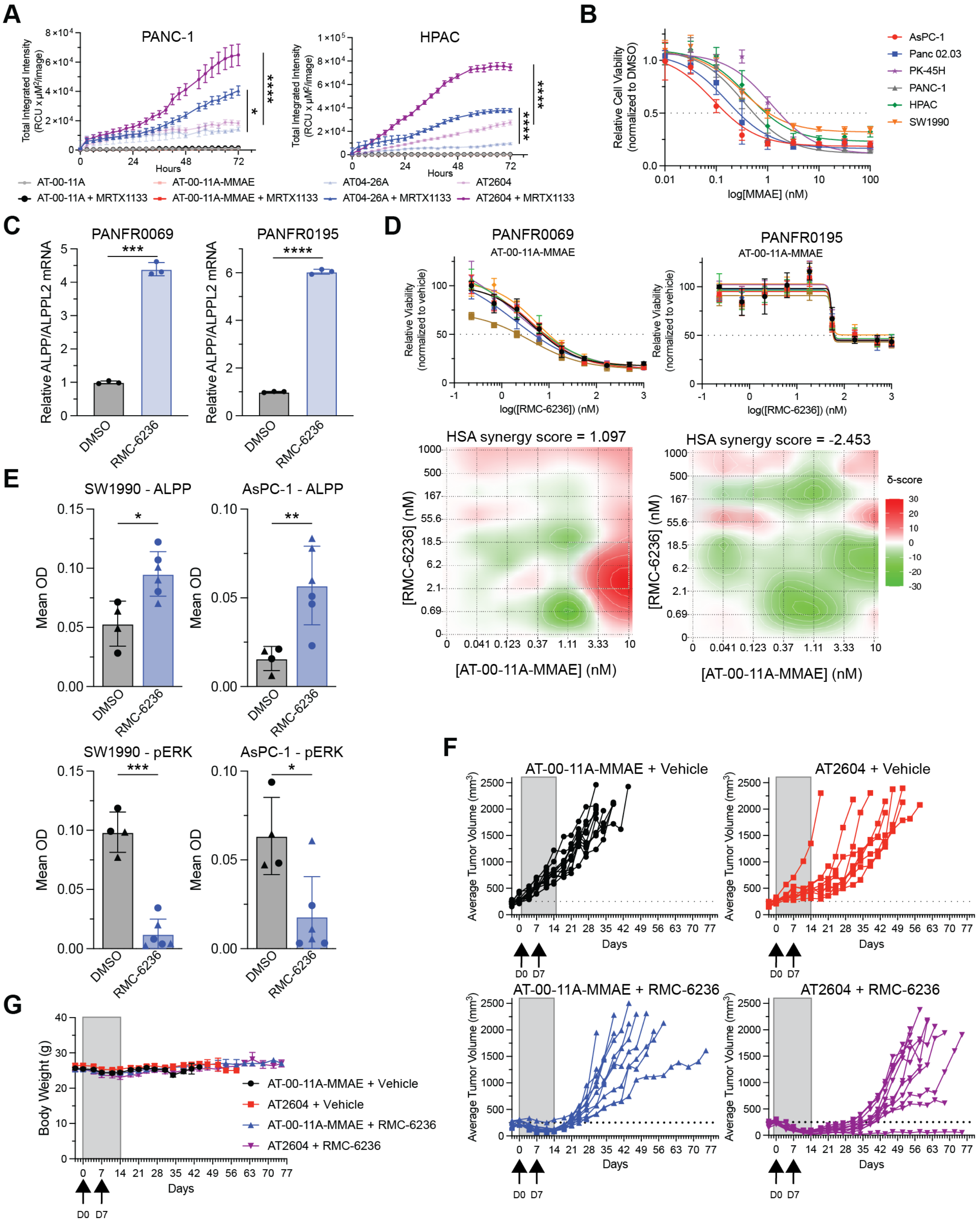
